# Spatiotemporal differences in salmon nutrient inputs restructure functional and taxonomic fungal communities in riparian system

**DOI:** 10.64898/2026.06.23.734103

**Authors:** Anne Y. Polyakov, Allen Larocque, Erik Lilleskov, Korena K. Mafune, Kristiina A. Vogt, Daniel J. Vogt, Andrew M. Berdahl

## Abstract

- Spawning salmon transport marine-derived nutrients (MDN) into riparian forests, influencing soil, plant, and animal communities, yet their effects on fungal communities remain poorly understood.
- We used DNA metabarcoding to examine fungal responses to three spatial patterns of salmon-derived nitrogen (N) in southwest Alaska: (i) patchy inputs from wildlife-deposited carcasses, (ii) a 21-year carcass relocation experiment, and (iii) natural N gradients with distance from streams.
- Decomposing carcasses increased saprotrophic fungal diversity, identifying taxa responsible for salmon carcass decomposition. Long-term carcass relocation reduced diversity of medium-distance fringe ectomycorrhizal fungi (EMF), whereas recent, patchy carcass inputs increased diversity of both medium-distance fringe and long-distance EMF—guilds often associated with low-nutrient environments. Along natural stream N gradients, EMF responses varied markedly within functional guilds and genera, revealing unexpected variation in N sensitivity among closely related taxa.
- Pulsed, spatially heterogeneous nutrient inputs enhanced diversity of typically nitrophobic EMF, likely reflecting their capacity to maintain extensive mycelial networks, exploit nutrient hotspots, and mobilize organic N and phosphorus. The diversity of responses along natural N gradients suggests that mechanisms linking EMF traits to nutrient acquisition and tolerance remain unresolved. Our findings emphasize the importance of linking fungal community composition with functional attributes and nutrient dynamics.

## Introduction

Growth-limiting resources are essential drivers of ecosystem structure and function because they limit primary productive capacity (Klock et al. 2022). Within riparian ecosystems, the terrestrial-aquatic interface is a fundamental pathway for nutrient exchange, notably illustrated by the migration of Pacific salmon (*Oncorhynchus* spp.) from marine to freshwater environments, which injects significant quantities of marine-derived nutrients (MDN) into riparian forests. These subsidies are incorporated into local food webs through animal consumption and microbial decomposition of carcasses (Holtgrieve et al. 2010, Minakawa et al. 2002, Field and Reynolds 2013, Larocque 2022). MDN in salmon carcasses include potentially important growth-limiting nutrients including nitrogen (N), phosphorus (P), magnesium (Mg), and calcium (Ca), which can significantly alter plant and animal community structure (Drake et al. 2005, Larocque and Simard 2023, Wagner and Reynolds 2019). For example, MDN can increase plant growth (Quinn et al. 2018) and reproductive allocation (Siemens et al. 2020), reduce plant diversity, and alter plant composition to increasingly nitrophilic and phylogenetically dispersed communities (Hocking and Reynolds 2012, Hurteau et al. 2016).

Despite the essential roles fungi play in forest ecosystems, the impact of MDN on fungal community structure, function, and implications for soil carbon (C) and nutrient dynamics remains poorly understood. Saprotrophic fungi are essential to the breakdown of complex organic compounds, which increases nutrient availability to plants (Northrup et al. 1995, Averill et al. 2014, Read et al. 2004), whereas mycorrhizal fungi form mutualisms with plant roots which benefits plants via increased nutrient acquisition in exchange for plant photosynthates, critical in improving tree nutrition and soil C storage (Averill et al. 2016, Hawkins et al. 2023). Therefore, the potential of fungi to mediate the long-term effects of MDN on plant communities is important to understand. In the only study to examine the effects of MDN on fungal communities, fungal species richness and diversity increased with higher salmon densities (Larocque 2022).

Fungi respond rapidly to environmental fluctuations and gradients, and are especially sensitive to changes in important limiting nutrients such as N (Zhao et al. 2020). Utilizing functional classifications—such as feeding strategies (saprotrophic vs. mycorrhizal) or hyphal exploration traits (Hobbie et al. 2014)—enables the testing of trait-based hypotheses (Nguyen et al. 2016, Zanne et al. 2020, Moeller et al. 2014). While saprotrophic fungi typically increase in relative abundance with increasing soil nutrient availability (Geng et al. 2023), long-term N addition has been shown to decrease N exchange between mycorrhizal fungi and plant hosts (Jach-Smith et al. 2020) and reduce mycorrhizal fungal biomass, richness, and diversity (Fierer 2017, Lilleskov et al. 2011, Treseder 2008). Although nutrient availability is a strong driver of mycorrhizal fungal community composition (Moeller et al. 2014), effects vary strongly by functional groups (Lilleskov et al. 2024). For example, in N- or P-limited environments, plant hosts favor ectomycorrhizal fungi (EMF) with long-distance exploration types, which use long-distance transport structures and large amounts of extraradical hyphae to forage over long distances and exploit nutrient patches (Moeller et al. 2014). Furthermore, these long-distance taxa also have strong enzymatic capabilities to break down and acquire organic N, increasing access to N pools (Hobbie and Agerer 2010). Similar to long-distance types, medium-distance fringe EMF exhibit high hyphal density and capacity for organic N use. In contrast, medium-distance smooth EMF feature hydrophilic hyphae with less extensive rhizomorphic structures and limited organic N mining capabilities (Lilleskov et al. 2011), and are selectively dominant with increasing nutrient availability (Moeller et al. 2014). Short-distance and contact EMF typically forage over small areas, have less C intensive exploration strategies, and are favored under high nutrient environments (Nara 2015).

Although altering soil N availability has direct consequences for EMF community composition and relative abundance, the specific mechanisms underlying these effects remain elusive (Lilleskov et al. 2019), partly because functional groups are not yet fully characterized (Hobbie et al. 2019). For example, mycorrhizal fungal response to elevated nutrients is not uniform, varying significantly both between and within functional groups. For instance, while high-N conditions typically cause long-distance, nitrophobic species like *Suillus bovinus* to decline, long-distance, nitrophilic species such as *Paxillus involutus* increase in abundance, as they appear to be specialized to acquire P under high-N, low-P conditions (Lilleskov et al. 2024). This indicates that additional functional traits must be considered to fully understand responses to nutrient availability.

Furthermore, the spatial and temporal distribution of nutrients influences fungal community composition. Increasing nutrient availability typically reduces the richness and abundance of long- and medium-distance foraging EMF due to decreasing belowground C flux from host plants (Lilleskov et al. 2011, Kutorga et al. 2013, Jach-Smith et al. 2020). However, in systems where nutrients are available but patchily distributed (i.e., hotspots) or where nutrient input occurs in pulses such as riparian zones along salmon streams, these functional groups might increase in relative abundance and richness due to their ability to maintain large network structures and mobilize organic N, making them more efficient at mining nutrient hotspots than plant roots (Moeller and Neubert 2016).

Streams with recurring annual spawning salmon typically show a decreasing N gradient away from the stream edge due to continual annual input of MDN over millennia (Feddern et al. 2019). This gradient is particularly evident in boreal forests (Feddern et al. 2019). Other factors also contribute to this N gradient, as N cycling generally decreases with distance from stream and with increase in slope due to changes in soil water content, and N fixed uphill will migrate down to the streams over time (Hogberg 1997, Kirchoff 2003).

This natural N gradient likely supports specific EMF communities adapted to varying soil N levels. However, research has not characterized how the abundance, diversity, and functional traits of these communities shift along this N gradient. Identifying the specific EMF functional groups adapted to higher soil N levels is crucial for understanding how these ecosystems will respond to anthropogenic N enrichment. Furthermore, while fungal communities associated with animal carcasses have been studied (Metcalf et al. 2015, Procopio et al. 2020), those specifically associated with salmon carcasses have not been characterized. Given the global decline in salmon populations—which threatens salmon-adapted fungal communities in riparian zones—this research aims to characterize the fungal taxa and functional roles near high-density salmon streams. This will provide a crucial reference system for identifying threatened, potentially vanishing EMF communities in regions with declining salmon populations, such as the Pacific Northwest.

This study investigated fungal community responses along three salmon-bearing streams in a southwest Alaskan boreal forest, focusing on the impacts of natural soil N gradients and salmon carcass decomposition (Figure 1). A key component of the research involved Hansen Creek, where a 21-year manipulation experiment (1997–2018) created a massive nutrient fertilization treatment by transferring over 200,000 kg of salmon carcasses from the stream and one bank to the other bank (Quinn et al. 2018). Our primary objectives were to evaluate how fungal community structure, relative abundance, richness, and diversity respond to: i) Short-term, localized nutrient pulses from individual, wildlife-deposited carcasses along stream banks; ii) Long-term, large-scale nutrient manipulation (the Hansen Creek experiment); and iii) Natural N gradients that occur with increasing distance from the stream edge.

**Figure 1.**
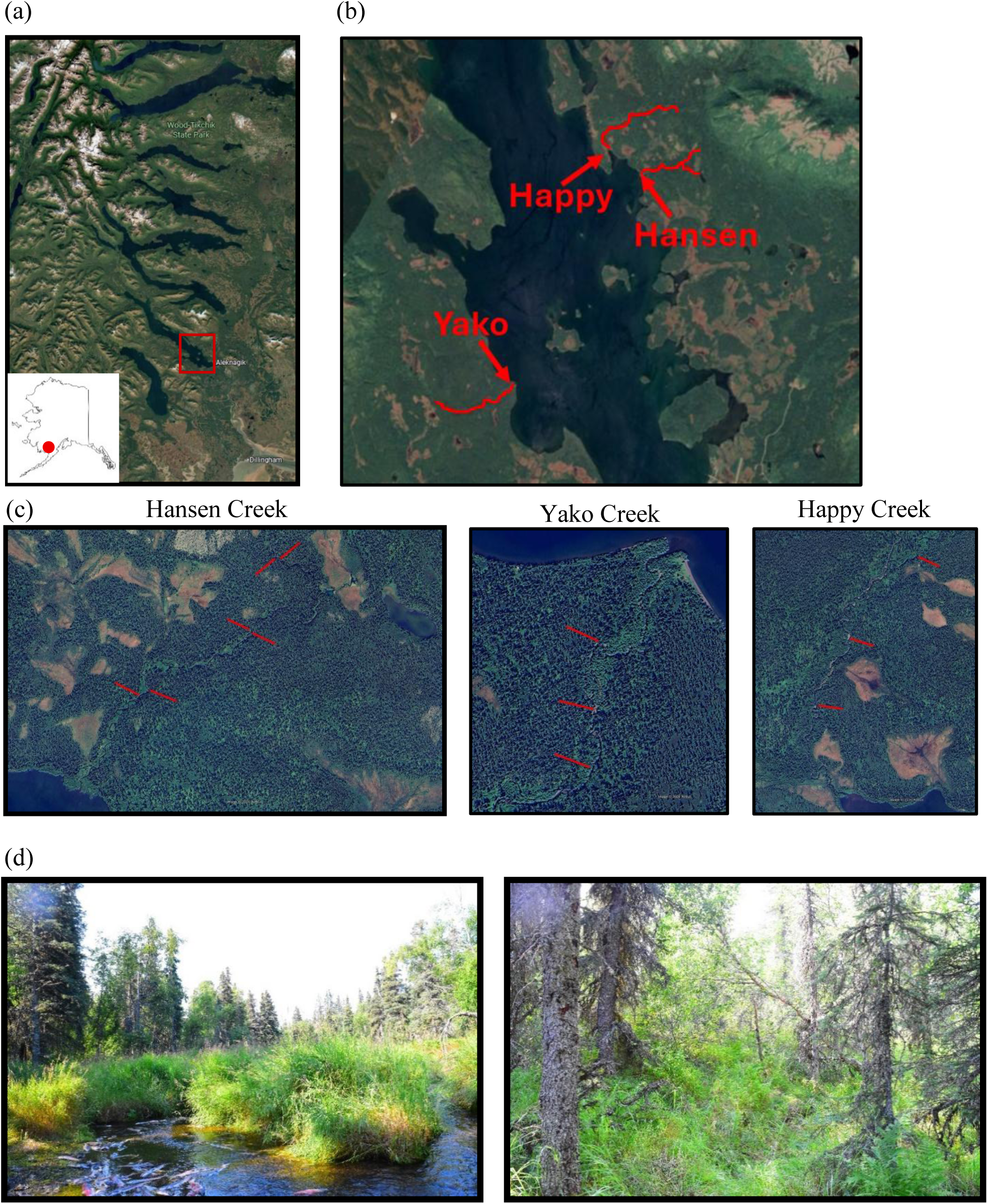
Study system. (a) The Wood River Lake system in southwest Alaska where (b) three salmon streams were sampled on Lake Aleknagik. (c) Transects along each of the three streams and (d) photographs of a study site at Hansen Creek close to the stream bank and 40 m into the forest.

We examined responses of fungal trophic guilds, EMF functional groups, and individual fungal genera separately. Since the carcass manipulation deposited considerably higher N than critical N loads associated with changes in fungal community structure (Lilleskov et al. 2024), we hypothesized that community composition would shift in response to the carcass manipulation, and that higher soil N levels from both MDN and along the N gradient would increase the relative abundance, richness, and diversity of saprotrophic fungi, but decrease those of long-distance and medium-distance fringe EMF.

## Materials and Methods

### Site description

This study was conducted at three salmon streams—Hansen, Happy, and Yako Creeks—that are tributaries to Lake Aleknagik within the Wood River System of southwest Alaska (Figure 1a). The surrounding boreal forest is dominated by white spruce (*Picea glauca*) and paper birch (*Betula papyrifera*). Riparian vegetation includes willow (*Salix* spp.), cottongrass (*Eriophorum* spp.), bracken fern (*Pteridium aquilinum*), horsetail (*Equisetum arvense*), and dwarf birch (*Betula nana*), with Vaccinium species, bryophytes, and heather (*Harrimanella stelleriana*) located further from the banks (Figure 1b). While green alder (*Alnus crispa*), a N-fixer that complicates marine-derived nutrient (MDN) tracing, was present at Happy Creek, it was absent from our sampling site at Hansen and Yako Creeks (Helfield and Naiman 2002). Soils are characterized primarily as well-drained Typic Haplocryods, formed in silty volcanic ash over gravelly glacial drift or loam till (Helfield and Naiman 2002). Hansen Creek was the site of a 21-year salmon carcass fertilization experiment, during which all salmon were removed from the stream and right bank, and thrown onto the left bank, depositing 8,000 kg of N and 1,356 kg of phosphorus (from 200,000 kg of salmon) on one side of the stream but not the other. Please see Quinn et al. (2018) and Feddern et al. (2019) for a complete description, with supporting details in the SI. Additionally, wildlife-deposited salmon carcasses were observed along all three study streams.

### Sample collection and processing

To characterize riparian soils, we established three sets of paired 100 m transects at Hansen Creek on opposing banks (six total, perpendicular to the stream) and three transects each at Happy and Yako Creeks. Transects targeted areas with representative vegetation—primarily white spruce (*Picea glauca*) and paper birch (*Betula papyrifera*)—and high annual salmon carcass input. At eight intervals along each transect (1, 3, 6, 10, 20, 40, 60, and 100 m from the active channel edge), we collected soil samples at 10–15 cm and 25–30 cm depths, targeting the deeper organic horizons (*Oe* and *Oa*) and the upper B (typically *Bhs*), respectively (USDA 2022). We determined gravimetric soil water content (g H_2_O/g dry soil) by weighing 50-100 g of field-moist soil, then drying at 105°C for 48 hours, and then reweighing (Gardner 1986). We froze all soil samples to −20 °C at the field site, transported them in a cooler to the University of Washington, and stored them there at −80°C until we extracted the DNA.

### Molecular and bioinformatics analysis

Genomic DNA was extracted from 5 mg of thawed moist soil using the DNeasy PowerSoil Pro Kit (Qiagen, Benelux BV) and quantified with a Qubit 4 Fluorometer using the dsDNA HS Assay Kit (Thermo Fisher Scientific, Wilmington, DE, USA). The fungal ITS1 region was targeted and amplified utilizing ITS1-F/ITS2 primers (White et al. 1990). Following the addition of Illumina adapters and polling to equimolar concentrations, sequencing was performed on the Illumina MiSeq platform (2 x 250 bp) at the University of Oregon Genomics Laboratory.

Raw reads were processed using CUTADAPT (v. X.X) to remove adapters (Martin 2011). Sequences were then analyzed with the DADA2 pipeline to filter (maximum expected errors: 2, minimum length: 500 bp) and dereplicate the reads, trim low-quality sequences, merge paired-ends, and remove chimeras (Callahan et al. 2016). The parameters maxEE = 4, 6 and truncLen = 270, 250, and minOverlap = 50 were used for ITS regions. Resulting Amplicon Sequence Variants (ASVs) were further clustered at 97% similarity using the DECIPHER package (v. 2.16.1) in R. Taxonomy was assigned via the naive Bayesian classifier (Wang et al. 2007) against the UNITE database (Nilsson et al. 2019). Community data were analyzed using the Phyloseq R package (McMurdie & Holmes 2013). Fungal trophic status and exploration types were assigned using FUNGuild (Nguyen et al. 2016), restricting analyses to “highly probable” or “probable” confidence rankings. For taxa with mixed trophic strategies (e.g., *Entoloma, Oidiodendron, Sistotrema, Cadophora*), trophic strategy was assigned at the species level (Brandrud et al. 2018).

### Statistical Analysis

To examine the effects of environmental variables on fungal community metrics, we implemented three sets of generalized linear mixed-effects models (GLMMs) (Table 1, S6). The first set analyzed the relative abundance, species richness, and Shannon diversity of saprotrophic and symbiotrophic trophic guilds. The second set examined these same metrics for ectomycorrhizal (EMF) functional guilds (short-distance, medium-distance fringe, medium-distance smooth, and long-distance). The third set modeled the relative abundance of individual EMF genera. We used negative binomial GLMMs for species richness (count data) and linear mixed-effects models for relative abundance and Shannon’s diversity index.

**Table 1.** Fungal genera placed into exploration type categories (long-distance, medium-distance fringe, medium-distance smooth, and short-distance (grouped by short-distance delicate, short-distance coarse, and contact) types from soil samples sampled along three salmon streams in Alaska. A list of species for each genus can be found in Table S6. *This genera contains both mycorrhizal and saprotrophic species, and these designations refer to the mycorrhizal species within this genera.

| **Long distance** | **Medium distance fringe** | **Medium distance smooth** | **Short distance delicate** | **Short distance coarse** | **Contact** |
| --- | --- | --- | --- | --- | --- |
| *Xerocomus*<br>*Paxillus*<br>*Alpova*<br>*Boletus* | *Cortinarius*<br>*Amphinema*<br>*Piloderma*<br>*Sistotrema* *<br>*Tricholoma*<br>*Lyophyllum* * | *Hydnum*<br>*Tomentella*<br>*Pseudotomentella*<br>*Thelephora*<br>*Lactarius*<br>*Entoloma* *<br>*Tomentellopsis* | *Laccaria*<br>*Inocybe*<br>*Naucoria*<br>*Tylospora*<br>*Hymenogaster*<br>*Tuber*<br>*Leotia*<br>*Hebeloma*<br>*Sebacina*<br>*Alnicola*<br>*Pulvinula*<br>*Endogone* | *Trichophaea*<br>*Wilcoxina*<br>*Cenococcum*<br>*Genabea*<br>*Otidea*<br>*Geopora* | *Russula*<br>*Amanita*<br>*Hygrophorus*<br>*Clavulina*<br>*Helvellosebacina* |

Fixed effect covariates included organic and mineral soil C:N ratios, organic soil moisture (gravimetric water content), organic soil pH, carcass proximity (binary, < 1 m, based on Drake et al. 2005), presence of carcass manipulation (binary, based on river bank), presence of a slope (binary, slope characterized as >10^∘^), and distance from the stream bank. To incorporate plant community composition, we performed a multiple correspondence analysis on plant species presence data (tree, shrub, herb, moss) within a 5 m radius (25*π* m^2^); the first five vectors (representing 60% of variance) were included as predictors (Table S4, Figure S4). Transect was included as a random effect.

Species indicator analysis was used to identify indicator species associated with the carcass manipulation and the presence of a nearby decomposing carcass using the indicspecies package in R (Table S5; Cáceres and Legendre 2009). Threshold Indicator Taxa Analysis (TITAN) using the TITAN2 package in R (Baker and King 2010) was implemented to identify the responses of individual fungal taxa to changes in organic soil C:N ratio (Figure 4). To examine the effect of the carcass manipulation on fungal community structure, the matrices of the pairwise taxonomic distance (Bray-Curtis) were calculated. Correlations between fungal community composition and environmental variables were evaluated using Mantel tests and the Bray-Curtis dissimilarity metric using the R package Ecodist (Goslee and Dean 2007). The significance of environmental variables on fungal community composition was evaluated with analysis of variance using distance matrices within the adonis2 function in the R package vegan (Oksanen et al. 2016). Distance-based redundancy analysis (dbRDA) was conducted with forward selection of the explanatory variables.

## Results

### Overall fungal community composition

We identified a total of 8754 fungal ASVs. Of these, 5583 were identified to genus, 6217 to family, and 8045 to phylum (Table S6). The identified ASVs included 10 phyla, 154 families, and 308 genera. Dominant phyla consisted of Basidiomycota, Mortierellomycota, and Ascomycota, with an average relative abundance of 67%, 18%, and 11% across all samples. Importantly, *Ascomycota* can be undersampled in the ITS1 region due to group 1 introns (Bellemain et al. 2010). The top ten most abundant genera were *Mortierella* (19%), *Russula* (13%)*, Laccaria* (9%)*, Inocybe* (8%)*, Tylospora* (7%)*, Cortinarius* (6%)*, Solicoccozyma* (5%)*, Lactarius* (3%)*, Tomentella* (2%), and *Naucoria* (2%), which constituted 84% of all sequences, indicating dominance of symbiotrophs (specifically EMF), relative to saprotrophic fungi (45% and 9% of total abundance, respectively). Within EMF, short-distance exploration types dominated (42% of total abundance).

### Effects of large-scale carcass manipulation on fungal communities

Fungal community composition significantly differed between the banks of Hansen Creek due to the carcass experiment (*p* = 0.05, R^2^ = 0.13, Figure S1). The carcass manipulation specifically reduced the relative abundance and species richness of medium-distance fringe EMF (Table 1, Table S6, Figure 2a). When examining the effect of carcass input by genera, the carcass manipulation increased the relative abundance of *Boletus, Tomentella, Thelephora, Tylospora, Meliniomyces,* and *Alnicola,* and decreased the relative abundance of *Cortinarius* and *Clavulina.* Species indicator analysis identified seven species associated with the carcass manipulation (Table S5a).

**Figure 2.**
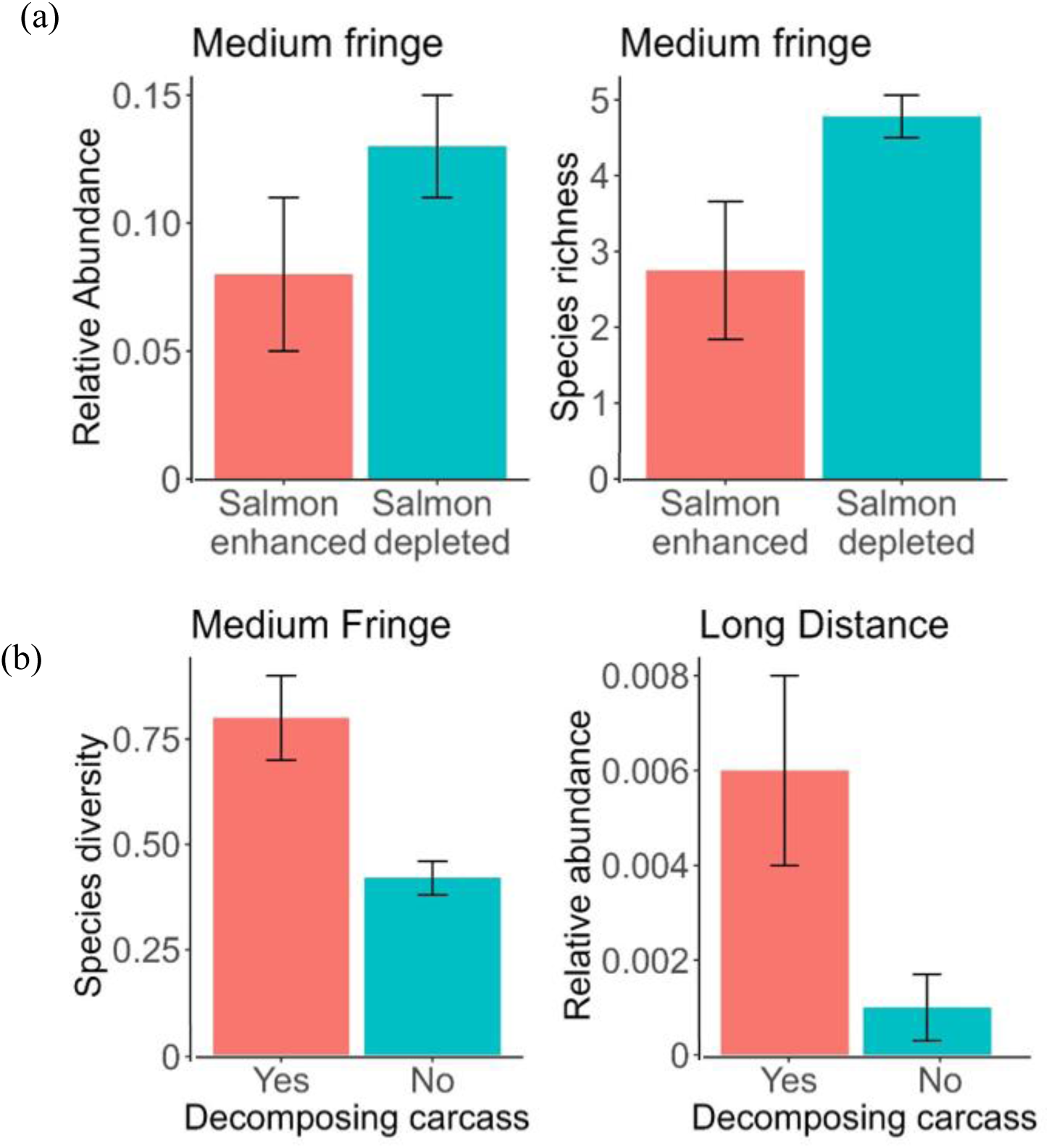
Predictions from the linear mixed effects models for (a) fractional relative abundance and species richness of medium-distance fringe fungi on the salmon-enhanced and salmon-depleted banks of Hansen Creek following a 21-year salmon carcass manipulation experiment, and (b) Shannon diversity index of medium-distance fringe fungi and fractional relative abundance of long-distance fungi at locations <1 m from a decomposing salmon carcass along three salmon streams in SW Alaska.

### Effects of local carcass proximity on fungal communities

Local carcass proximity significantly altered fungal community composition (*p* = 0.03, R^2^ = 0.15). Decomposing carcasses increased both the species diversity of medium-distance fringe and the relative abundance of long-distance EMF exploration types (Figure 2b). At the genus level, proximity to carcasses led to higher relative abundances of *Paxillus*, *Xerocomus*, *Piloderma*, *Sistotrema*, *Hydnum*, *Geopora*, and *Helvellosebacina*, while *Trichophaea* and *Hebeloma* decreased. Species indicator analysis identified ten species associated with the presence of a nearby decomposing carcass (Table S5a). Local decomposing carcasses also increased the richness and diversity of saprotrophic fungi (Figure 3), and we identified 35 genera (Table 2) and 51 species of saprotrophic fungi that were found exclusively near decomposing carcasses, including multiple saprotrophic *Entoloma* and *Mortierella* species (Table S1, Table S5a).

**Figure 3.**
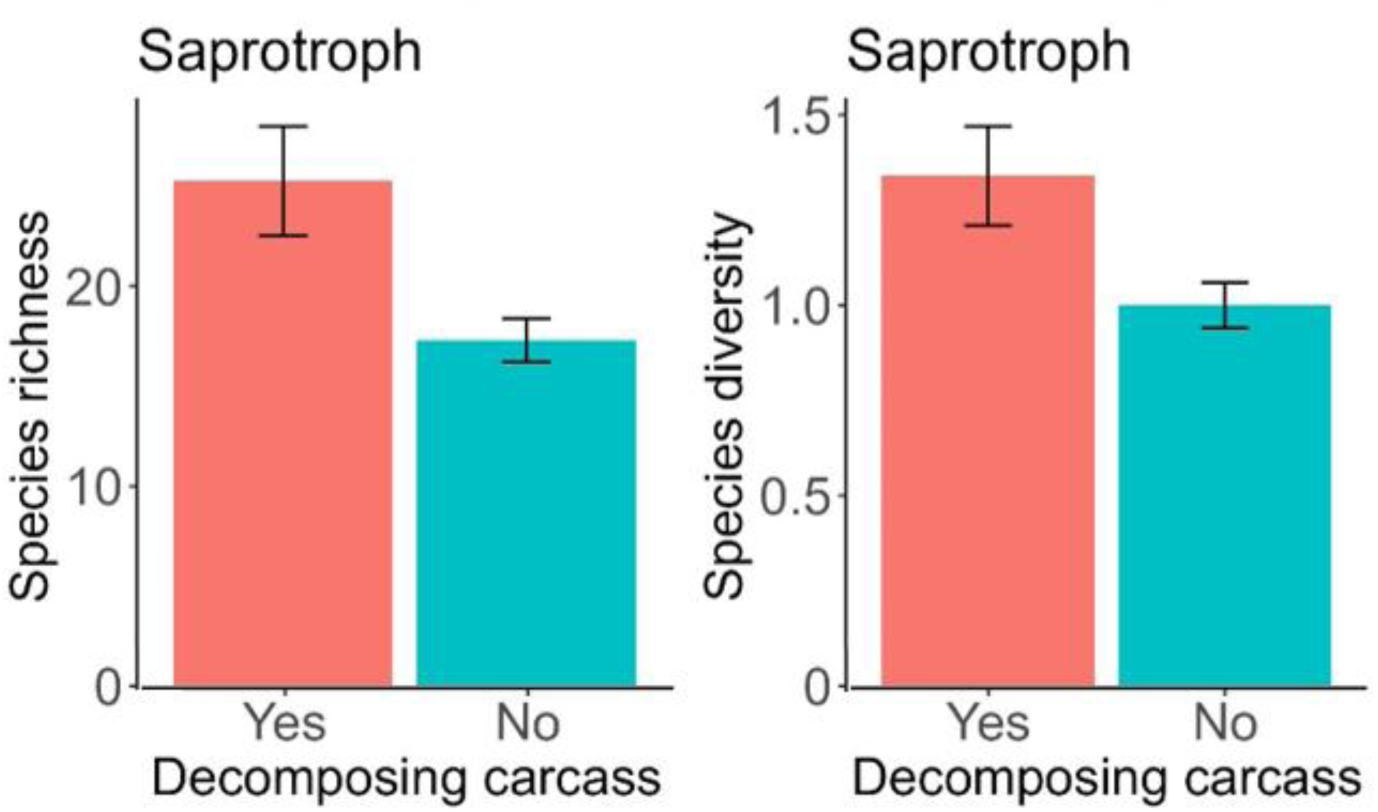
Predictions from the linear mixed effect models for species richness and Shannon diversity index of saprotroph fungi at locations <1 m from a decomposing salmon carcass along three salmon streams in SW Alaska.

**Table 2.**
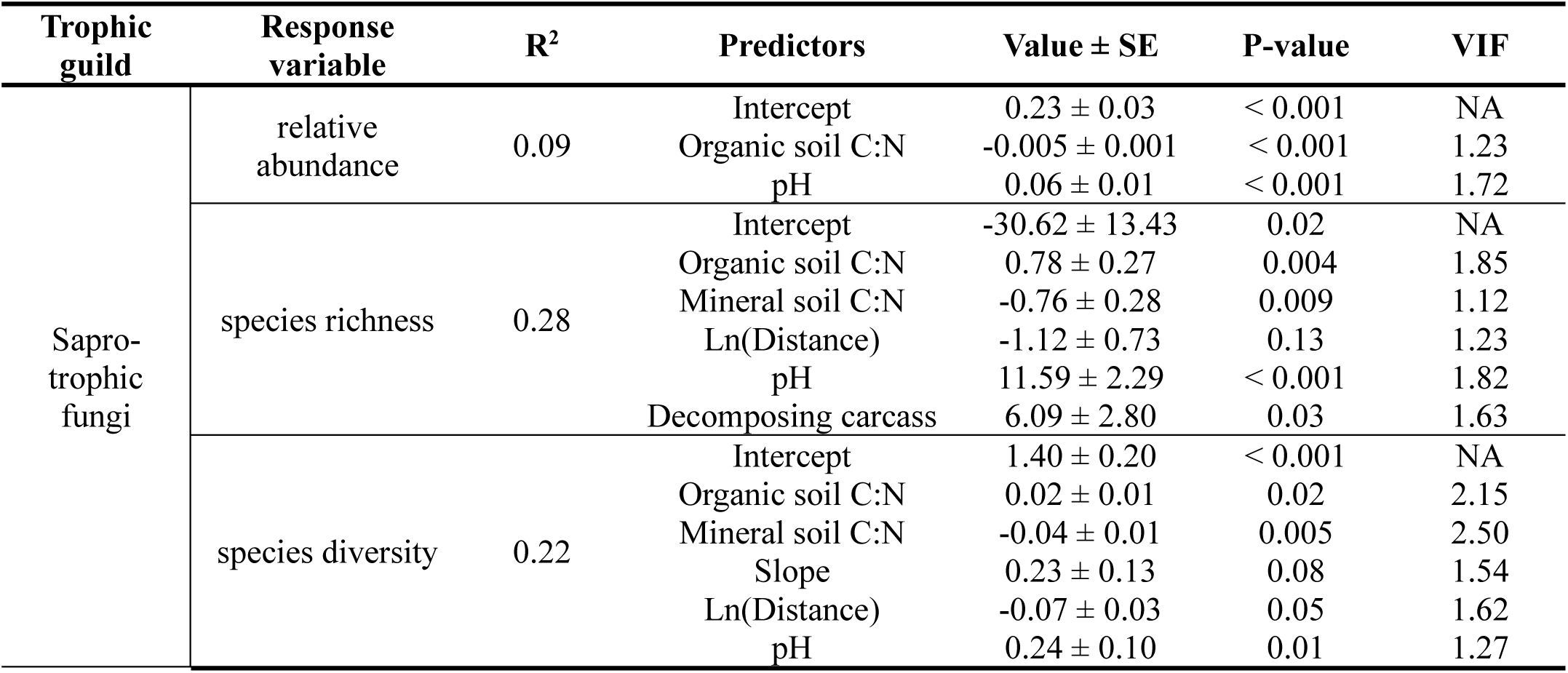
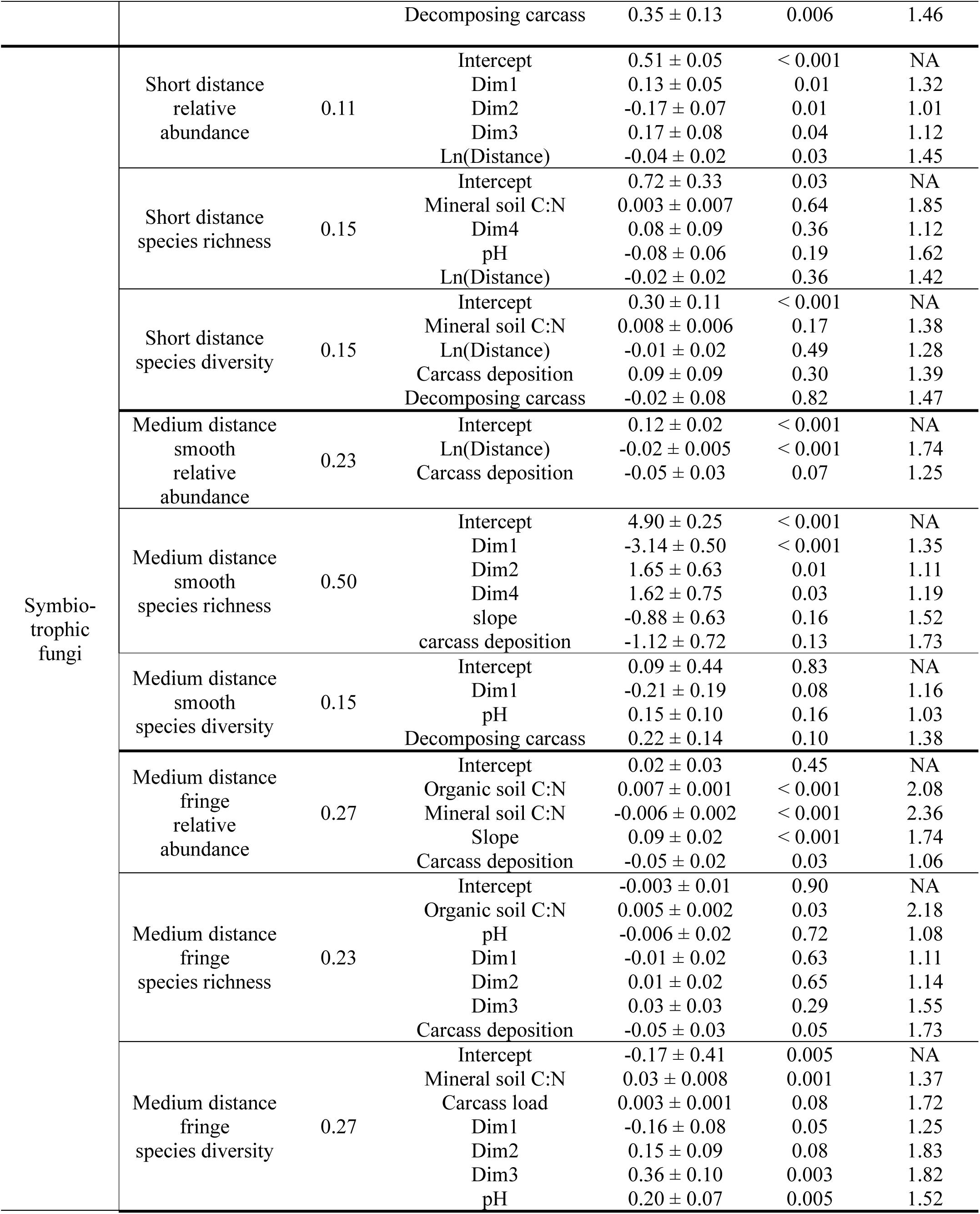
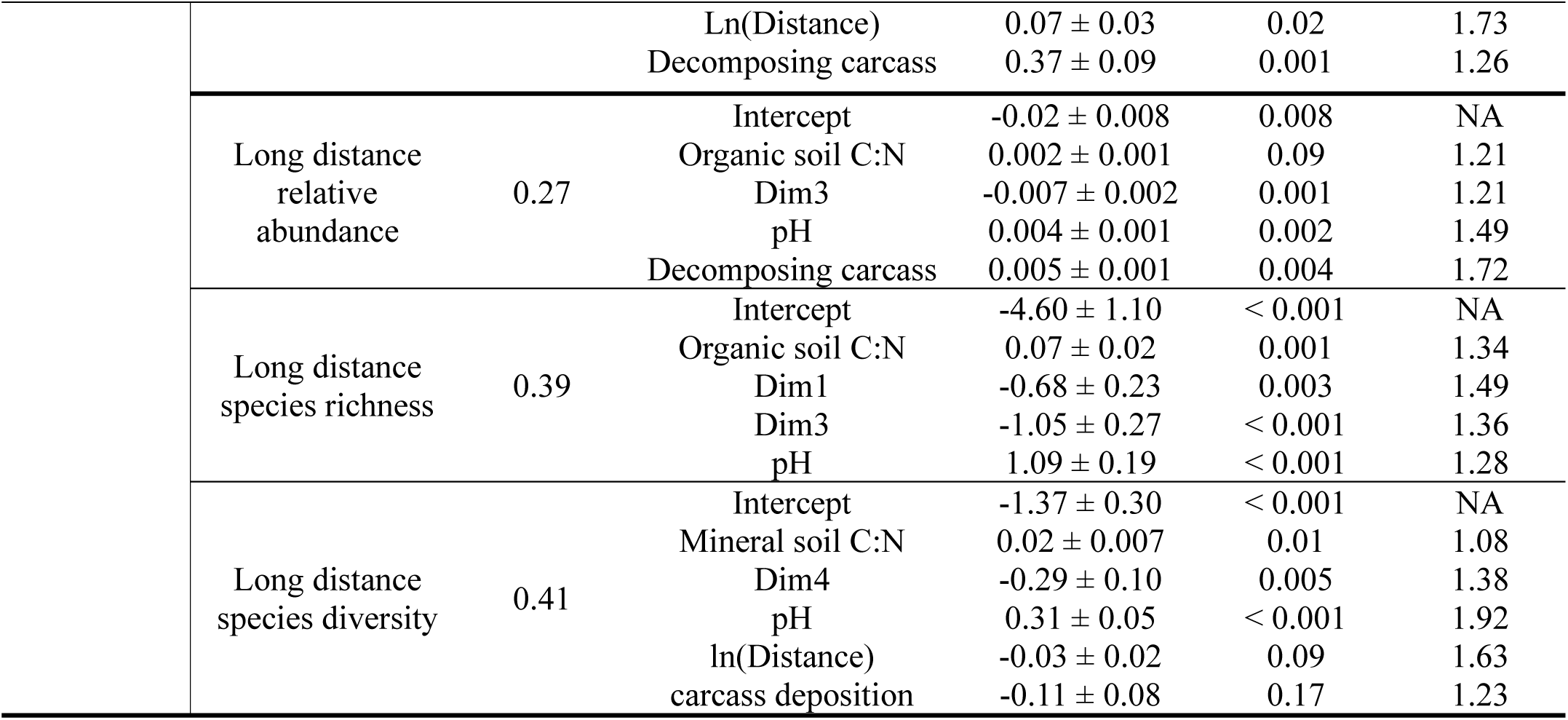
Predictors from the linear mixed effect models for relative abundance, species richness, and species diversity of saprotrophic and symbiotrophic (long-distance, medium-distance fringe, medium-distance smooth, and short-distance) fungi along three salmon streams in SW Alaska. Models include conditional R^2^, all predictor variables, beta coefficient estimates ± SE, P values and VIF (variance inflation factor). All models included a random effect of transect nested within stream. GWC indicates gravimetric water content. Dim1-Dim4 describe plant community composition based on multiple correspondence analysis (Figure S4).

*Fungal communities along soil N gradients* - Fungal community dynamics shifted significantly across N gradients, with richness and diversity of both saprotrophs and symbiotrophs correlating positively with organic soil C:N ratios and negatively with mineral soil C:N ratios (Table 2). Within the symbiotrophs, medium-distance fringe ectomycorrhizal fungi (EMF) showed increases in both relative abundance and richness as organic soil C:N increased, while long-distance EMF increased only in abundance (Table 2).

TITAN analysis revealed substantial variation in strategies within functional groups and genera. While many EMF species responded to N availability, these responses were often species-specific (Figure 4). For *Cortinarius*, most species favored higher organic C:N ratios, though *C. caninus* declined. For *Russula*, *R. favrei* associated with lower C:N ratios, whereas *R. suecica*, *R. paludosa*, *R. consobrina*, and *R. claroflava* thrived at higher ratios. For *Lactarius*, *L. rufus* and *L. atroviridis* increased with higher C:N ratios, while *L. necator* and *L. glyciosmus* decreased. For *Entoloma*, both symbiotrophic and saprotrophic species generally increased under higher N availability (lower C:N). For long-distance EMF, despite general trends, specific types like *Paxillus involutus* and *Alpova corsicus* decreased as organic soil C:N ratios rose.

### Nutrient limitation in white spruce and paper birch

Nutrient limitation analyses revealed no significant difference in N:P ratios for white spruce (*p*=0.3) or paper birch (*p*=0.5) between salmon-enhanced and salmon-depleted banks (1–6 m, Figure S2, Table S3). White spruce foliar chemistry (mean 1-6 m N:P: 7.3 ± 1.1; 1-100 m %N: 1.06 ± 0.14) indicated N limitation, while %P (0.15 ± 0.02) suggested sufficient P (Carter 1992, Güsewell et al. 2004; Figure S2, Table S3). Conversely, paper birch (mean 1-6 m N:P: 9.8 ± 2.3) suggested potential N and P co-limitation, with 1-100 m N concentrations (2.18 ± 0.31%) indicating slight N limitation (Keski-Saari and Julkunen-Tiitto 2003, Wang et al. 1998; Figure S2, Table S3).

## Discussion

### Long-term carcass manipulation suppresses N-sensitive EMF

Consistent with our hypotheses, fungal community composition differed significantly between the salmon-enhanced and salmon-depleted stream banks at Hansen Creek following long-term carcass addition (Figure S1), showing declines in relative abundance and species richness of medium-distance fringe EMF on the salmon-enhanced bank (Table 1, Table S6, Figure 2a). Medium-distance fringe EMF are among the most N-sensitive groups and are likely affected because they utilize carbon-intensive foraging strategies to mobilize organic N which are disfavored when inorganic N is abundant (Agerer 2001, Lilleskov et al. 2011, Lindahl and Tunlid 2015).

When inorganic N is abundantly added by salmon carcasses, host plants reduce carbon allocation to these costly partners in favor of lower-cost symbionts or direct root nutrient uptake (Jach-Smith et al. 2020, Lilleskov et al. 2024). For example, many species within the genus *Cortinarius*, a lineage widely recognized as nitrophobic (Bödeker et al. 2014, Lilleskov et al. 2024), declined in relative abundance in response to long-term carcass addition (Table S2). The shifts in the EMF community persisted even two years after the carcass addition ceased, indicating long recovery times for these communities following sustained, high-magnitude nutrient inputs.

### Patchy, short-term carcass inputs favor network-forming EMF

In contrast to the effects of chronic enrichment, naturally occurring decomposing carcasses increased relative abundance of long-distance and diversity of medium-distance fringe EMF in their immediate vicinity (Figure 2b, Table S1, Table S2, Table S5a). Notably, several taxa (especially from *Cortinarius*) that had declined under long-term carcass manipulation increased near individual decomposing carcasses, indicating that the same fungal lineages can respond in opposite directions depending on the spatial and temporal scale of nutrient delivery. Long-distance and medium-distance fringe EMF possess extensive extraradical mycelial networks, rhizomorphs, and hydrophobic hyphae that facilitate efficient exploitation of spatially discrete nutrient patches (Moeller et al. 2014, Lilleskov et al. 2024). These traits likely confer a competitive advantage in environments characterized by pulsed, heterogeneous nutrient inputs such as salmon carcasses deposited by wildlife (Moeller and Neubert 2016). Foliar nutrient analyses indicated N limitation in white spruce and potential N–P co-limitation in paper birch (Figure 3, Figure S4), suggesting that host plants may increase carbon allocation to network-forming fungi to efficiently access nutrients from nearby carcasses. This was also supported by the presence of fungal species specialized for both N uptake (some *Cortinarius* species) and P uptake (e.g., *Paxillus involutus* and *Alpova corsicus*; Lambers et al. 2006, Lilleskov et al. 2024, Plassard et al. 2011). Because salmon carcasses contain multiple essential nutrients, including P, magnesium, and calcium (Drake et al. 2005, Larocque and Simard 2023, Wagner and Reynolds 2019), these fungi may enhance host access to a broad suite of limiting resources rather than N alone. Together, these patterns suggest that patchy nutrient hotspots favor EMF with extensive mycelial networks and flexible nutrient acquisition strategies.

Decomposing salmon carcasses also increased the richness and diversity of saprotrophic fungi (Figure 3). While fungal communities associated with vertebrate carcasses have been documented in other systems (Metcalf et a. 2015, Procopio et al. 2020, Sagara et al. 2008, Sagara 1976), saprotrophic fungi associated with salmon carcasses have not previously been characterized using high-throughput sequencing. The occurrence of numerous saprotrophic taxa (Table 2) exclusively near decomposing carcasses suggests that salmon inputs support a distinct decomposer community (Table S1, Table S5a) that may contribute to rapid organic matter breakdown and nutrient release in riparian soils.

### Fungal responses along a natural soil C:N gradient

Fungal community richness—both saprotrophic and symbiotrophic—increased as organic soil C:N ratios rose but dropped as mineral soil C:N ratios increased. Based on the peaks of cumulative distributions of change points for fungal species, those responding negatively to organic soil C:N were most often found at a ratio < 19 (N is not limiting) while species responding positively were most often found at ratios > 25 (N is limiting, Figure 5; Brust 2019). These thresholds define key structural changes in the fungal community within this system. Among EMF, medium-distance fringe species showed gains in both richness and relative abundance in response to organic C:N ratios, while long-distance EMF increased only in relative abundance (Table 2). However, TITAN analysis showed that species-level responses varied widely within functional groups and genera, indicating substantial interspecific variation in nutrient use strategies (Figure 4).

**Figure 4.**
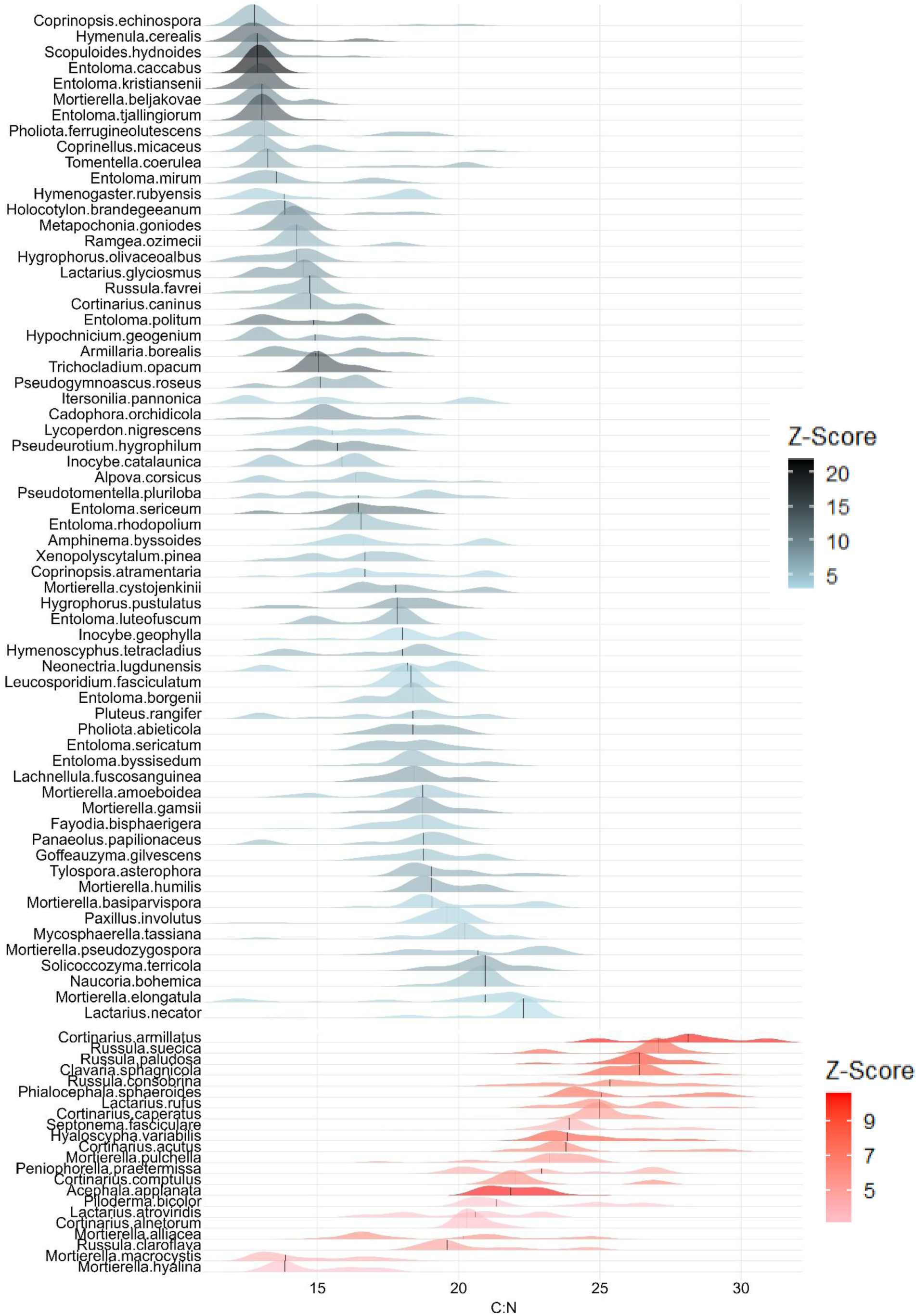
TITAN analysis shows the probability densities of change points for fungal species with negative (blue) and positive (red) responses to increasing organic soil C:N along three salmon streams in SW Alaska.

**Figure 5.**
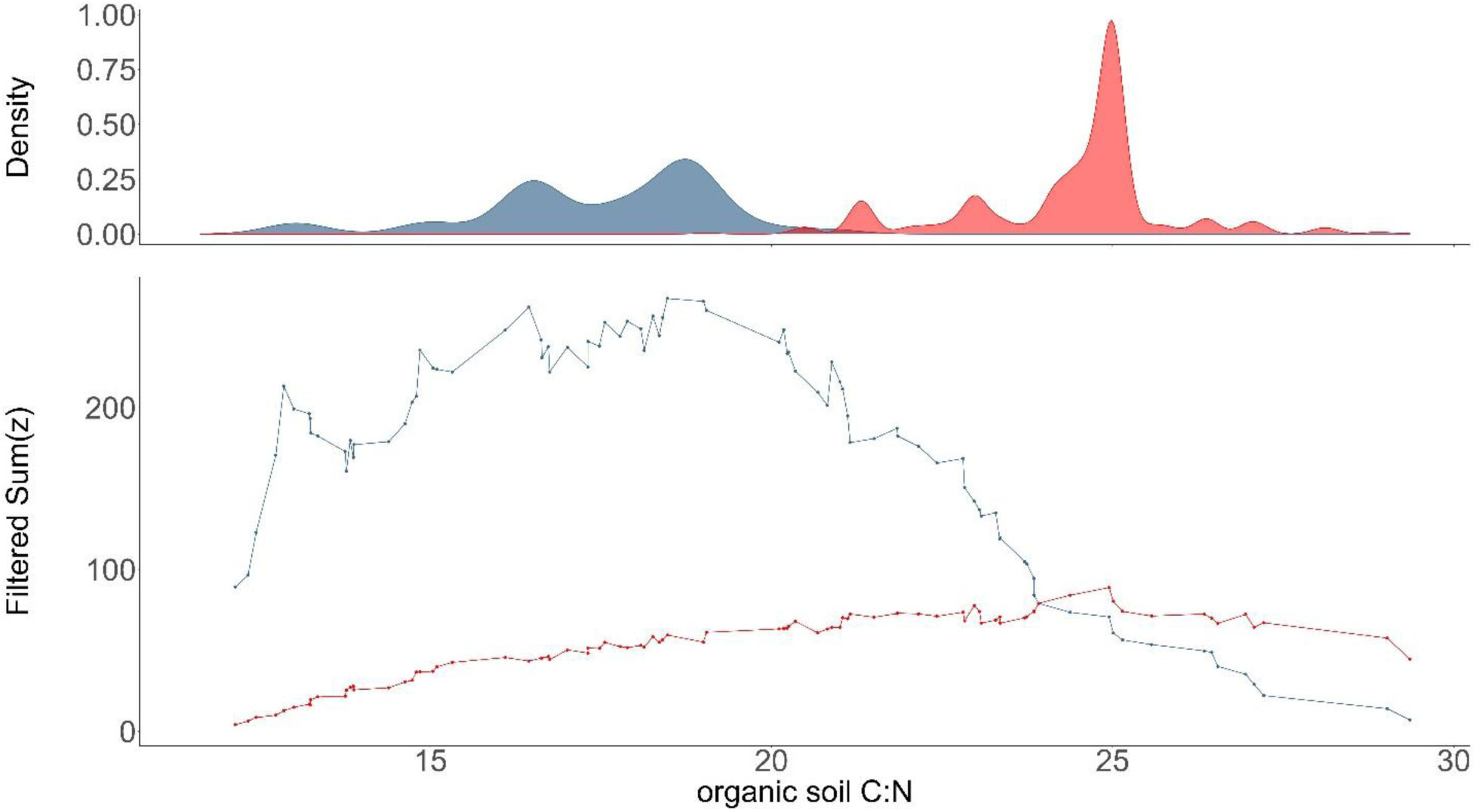
The top panel shows probability densities of change points for decreases (blue) and increases (red) as determined from bootstrap replicates, and the bottom panel shows the sum of z-scores for fungal species with negative (blue) and positive (red) z-scores. Peaks represent fungal community thresholds for change along organic soil C:N gradient along three salmon streams in SW Alaska.

For example, although most *Cortinarius* species increased under N-limited conditions (Figure 4, Table S2), *Cortinarius caninus* responded positively to higher N availability, suggesting functional differentiation within this genus. *Cortinarius* is recognized as a highly N-sensitive group, likely due to its oxidative enzymes (e.g., peroxidases) that can mobilize organically-bound N (Bödeker et al. 2014, Lilleskov et al. 2019). The sensitivity of *Cortinarius* to N fertilization may result from the selection against EMF capable of mobilizing organic N under high N conditions, though it remains unclear whether this selection is driven by host allocation, host selection, or soil-mediated direct effects (Lilleskov et al. 2024). Our results suggest that *Cortinarius caninus* could potentially have fewer peroxidase copy numbers or lower peroxidase activity than other *Cortinarius* species, but further functional studies are needed. Similar variability in N-response among *Russula* and *Lactarius* species challenges the classification of these genera as uniformly nitrophilic or nitrophobic, emphasizing that nutrient acquisition traits are highly variable at the species level.

Our findings indicate that *Russula* species exhibit diverse, widespread sensitivity to soil C:N ratios and, consequently, N availability (Lilleskov et al. 2011, Van der Linde et al. 2018), While *Russula favrei* showed higher relative abundance at lower soil C:N ratios (high N), other species—*Russula suecica*, *R. paludosa*, *R. consobrina*, and *R. claroflava*—increased in abundance with higher C:N ratios, signaling N sensitivity. Some *Russula* species have retained peroxidase genes (Looney et al. 2022), and these four species could be candidates for some retention of peroxidase genes due to their response to N. Surprisingly, although *Russula consobrina* is a member of a nitrophilic group of *Russula* (Avis 2012) and was associated with sites with decomposing carcasses (Table S5a), the species decreased with higher soil N levels along a gradient (Figure 4). This suggests opposing, context-dependent effects of high N from carcass decomposition versus soil N gradients on this species.

Although *Lactarius* is generally considered nitrophilic (Moeller et al. 2015, Lilleskov et al. 2019), our results reveal varied responses to elevated N. *Lactarius rufus* and *L. atroviridis* increased with higher soil C:N ratio (N-sensitive), whereas *Lactarius necator* and *L. glyciosmus* decreased with higher C:N ratio (N-tolerant). Interestingly, *Lactarius rufus* was strongly associated with carcass sites (Table S5a), similar to *Russula consobrina*, although *Lactarius rufus* varies in its response to N (Lilleskov et al. 2011).

*Entoloma* species, including both saprotrophic and symbiotrophic types, generally increased with higher soil N, indicating potential N-tolerance for the genus (Table 4). The very high N concentrations in sporocarps of *Entoloma* (roughly twice that of co-occurring genera) (Chen et al. 2019) suggests that its N demand must be quite high. Overall, multiple EMF species increased in relative abundance with increasing soil N (Figure 4), indicating that these species might be adapted to respond positively to anthropogenic N enrichment.

The relative abundance of long-distance EMF types *Paxillus involutus* and *Alpova corsicus* decreased with organic soil C:N ratio (Figure 4). Although some groups of long-distance EMF such as *Suillus* species are highly sensitive to N deposition, others such as *Paxillus involutus* and *Alpova corsicus* are nitrophilic species that increase with higher soil N levels (Lilleskov et al. 2024) and are associated with P limitation. *Alpova corsicus* is highly associated with N-fixing plant hosts, and likely specializes in other resources such as P (Rochet et al. 2011). *Paxillus involutus* is hypothesized to be specialized for P acquisition in high N, low P environments (Lilleskov et al. 2024). It is likely that distinct clades of long-distance EMF are associated with either N limitation or P limitation, but not both, and are specialized for either P uptake under P limitation or N uptake under N limitation. This exploration type could be associated with high belowground C allocation when aboveground growth is limited by either N or P (Lilleskov et al. 2024).

Furthermore, there may be a link between high N and alder-associated communities, as both *Alpova corsicus* and *Paxillus involutus* are associated with N-fixing alders (Walker et al. 2014), and the presence of other alder-associated generas such as *Naucoria* further suggests this. In fact, some species of nitrophilic *Cortinarius* and *Tomentella* are also alder associates (Walker et al. 2014). However, *Cortinarius caninuş*which responded positively to higher N at these sites, has no evidence of associating with alder, and was not found at sites with alder. Furthermore, *Naucoria bohemica*, which also increased with higher soil N, was found at many sites without alder. Therefore, the high N communities at these sites are not restricted to alder communities.

### Spatiotemporal scale mediates fungal responses to salmon nutrients

Our results show that fungal community responses to salmon-derived nutrients are strongly mediated by the spatial and temporal dynamics of nutrient inputs. Chronic, high-magnitude nutrient enrichment produced responses resembling those observed under anthropogenic N deposition, including declines in N-sensitive EMF. In contrast, naturally deposited carcasses created small-scale, transient nutrient hotspots that favored network-forming fungi adapted to patchy resource availability. These findings suggest that fungal communities are shaped not only by nutrient availability, but also by how nutrients are distributed in space and time. Consistent with broader ecological theory (Lodge et al. 1994, Weber and Brown 2013), pulsed nutrient inputs may promote higher fungal diversity than continuous enrichment.

### Implications for ecosystem function and conservation

Understanding these fungal communities can also assist in the conservation of certain functional groups whose populations have been declining. Increased soil N availability due to anthropogenic factors such as N pollution is causing the global decline of functionally related EMF taxa with long-distance, as well as medium-distance fringe and medium-distance mat, exploration types (Lilleskov et al. 2011). The functional group’s severe decline or regional extinction could have significant consequences for forest ecosystems. By losing these fungal taxa, many forests will not only lose or severely decline in the ability to acquire N from organic N pools, but have a reduced capacity to sequester soil C (Averill et al. 2018), which may have further consequences for additional forest functions such as uptake of other nutrients, water uptake, and pathogen resistance.

Arnolds (2007) reported that ecosystems across Europe experiencing long-term input of high N deposition are those where thousands of fungal species are included in Red Lists (e.g., vulnerable to extinction). Also, a third of the species listed on the Red Lists are ectomycorrhizal fungi where regional extinctions have already been recorded for some fungi (Arnolds 2007, Arnolds 2010). However, these EMF communities are poorly characterized, and we still lack critical ecological knowledge of this important group of fungi (Lilleskov et al. 2019). DNA-based molecular tools can provide us with new insights concerning the diversity of these fungal communities to help us better assess the implications of losing them for fungal conservation biology and ecosystem function.

## Conclusions

The effects of MDN on soil fungal communities depends on the spatial and temporal scale of salmon inputs. A 20-year experiment simulating salmon nutrient deposition showed that high N levels selected against medium-distance fringe EMF that have high C demand and peroxidase activity, resulting in decreased relative abundance and species richness of this group. In contrast, individual decomposing salmon carcasses (placed by wildlife) increased the relative abundance of long-distance EMF and enhanced the species diversity of medium-distance fringe types, particularly *Cortinarius* species. These fungi likely thrive due to their ability to maintain large network structures, mine nutrient hotspots, and access organic N and P. Decomposing carcasses significantly increased the species richness and diversity of saprotrophic fungi, creating unique, localized communities, specifically identified in this study.

The study suggests that preserving salmon habitat and populations is crucial for maintaining riparian EMF and saprotrophic fungal diversity. Furthermore, using areas with high spawning salmon populations as a reference system can help identify and protect these potentially threatened fungal communities in regions with declining salmon runs. The varied responses of fungal species to N gradients indicate that the specific mechanisms underlying these shifts require further investigation.

## Acknowledgements

We would like to thank the Alaska Salmon Program for making it possible to perform this research. We would also like to thank the Puget Sound Mycological Society, the Oregon Mycological Society, the Stuntz Foundation, and the Mycological Society of America for helping fund this research.

## Author Contributions

Conceptualization, A.Y.P., G.H., A.L., E.L., K.V., K. K. M., D.V., and A.M.B.; Methodology, A.Y.P., G.H., A.L., E.L., K.V., K. K. M., D.V., and A.M.B.; Statistical Analysis, A.Y.P., G.H., E.L., A.M.B.; Investigation, A.Y.P., G.H., A.L., E.L., K. K. M., K.V., D.V., and A.M.B.; Writing & Editing, A.Y.P., G.H., A.L., E.L., K. K. M., K.V., D.V., and A.M.B.;

## Data Availability

All data and code have been archived at Harvard Dataverse (https://doi.org/10.7910/DVN/EMUQXF). All analyses are fully reproducible.

## References

Arnolds, E. J. M. (2007). Biogeography and conservation. In C. P. Kubicek & I. S. Druzhinina (Eds.), Environmental and microbial relationships (The Mycota IV, pp. 105–124). Springer.

Arnolds, E. (2010). The fate of hydnoid fungi in The Netherlands and northwestern Europe. Fungal Ecology, 3(2), 81–88.

Averill, C., & Hawkes, C. V. (2016). Ectomycorrhizal fungi slow soil carbon cycling. Ecology Letters, 19(8), 937–947.

Averill, C., Dietze, M. C., & Bhatnagar, J. M. (2018). Continental-scale nitrogen pollution is shifting forest mycorrhizal associations and soil carbon stocks. Global Change Biology, 24(10), 4544–4553.

Averill, C., Turner, B. L., & Finzi, A. C. (2014). Mycorrhiza-mediated competition between plants and decomposers drives soil carbon storage. Nature, 505(7484), 543–545.

Avis, P. G. (2012). Ectomycorrhizal iconoclasts: The ITS rDNA diversity and nitrophilic tendencies of fetid *Russula*. Mycologia, 104(5), 998–1007.

Baker, M. E., & King, R. S. (2010). A new method for detecting and interpreting biodiversity and ecological community thresholds. Methods in Ecology and Evolution, 1(1), 25–37.

Bellemain, E., Carlsen, T., Brochmann, C., Coissac, E., Taberlet, P., & Kauserud, H. (2010). ITS as an environmental DNA barcode for fungi: An in silico approach reveals potential PCR biases. BMC Microbiology, 10, 189.

Bödeker, I. T. M., Clemmensen, K. E., de Boer, W., Martin, F., Olson, Å., & Lindahl, B. D. (2014). Ectomycorrhizal *Cortinarius* species participate in enzymatic oxidation of humus in northern forest ecosystems. New Phytologist, 203(1), 245–256.

Brandrud, T. E., Bendiksen, E., Jordal, J. B., Weholt, Ø., Eidissen, S. E., Lorås, J. A., Dima, B., & Noordeloos, M. E. (2018). Entoloma species of the rhodopolioid clade (subgenus Entoloma; Tricholomatinae, Basidiomycota) in Norway.

Brust, G. E. (2019). Management strategies for organic vegetable fertility. In Safety and practice for organic food (pp. 193–212). Academic Press.

de Cáceres, M., & Legendre, P. (2009). Associations between species and groups of sites: Indices and statistical inference. Ecology, 90(12), 3566–3574.

Callahan, B. J., McMurdie, P. J., Rosen, M. J., Han, A. W., Johnson, A. J. A., & Holmes, S. P. (2016). DADA2: High-resolution sample inference from Illumina amplicon data. Nature Methods, 13(7), 581–583.

Carter, R. E. I. D. (1992). Diagnosis and interpretation of forest stand nutrient status. In Forest fertilization: Sustaining and improving nutrition and growth of western forests (College of Forest Resources Contribution 73, pp. 90–97).

Chen, J., Heikkinen, J., Hobbie, E. A., Rinne-Garmston, K. T., Penttilä, R., & Mäkipää, R. (2019). Strategies of carbon and nitrogen acquisition by saprotrophic and ectomycorrhizal fungi in Finnish boreal *Picea abies*-dominated forests. Fungal Biology, 123(6), 456–464.

Drake, D. C., Smith, J. V., & Naiman, R. J. (2005). Salmon decay and nutrient contributions to riparian forest soils. Northwest Science, 79, 61–71.

Feddern, M. L., Holtgrieve, G. W., Perakis, S. S., Hart, J., Ro, H., & Quinn, T. P. (2019). Riparian soil nitrogen cycling and isotopic enrichment in response to a long-term salmon carcass manipulation experiment. Ecosphere, 10(11), e02958.

Field, R. D., & Reynolds, J. D. (2013). Ecological links between salmon, large carnivore predation, and scavenging birds. Journal of Avian Biology, 44, 9–16.

Fierer, N. (2017). Embracing the unknown: Disentangling the complexities of the soil microbiome. Nature Reviews Microbiology, 15(10), 579–590.

Gardner, W. H. (1986). Water content. In A. Klute (Ed.), Methods of soil analysis: Part 1—Physical and mineralogical methods (pp. 493–544). Soil Science Society of America.

Geng, X., Zuo, J., Meng, Y., Zhuge, Y., Zhu, P., Wu, N., Bai, X., Ni, G., & Hou, Y. (2023). Changes in nitrogen and phosphorus availability driven by secondary succession in temperate forests shape soil fungal communities and function. Ecology and Evolution, 13(10), e10593.

Goslee, S. C., & Urban, D. L. (2007). The ecodist package for dissimilarity-based analysis of ecological data. Journal of Statistical Software, 22, 1–19.

Güsewell, S. (2004). N:P ratios in terrestrial plants: Variation and functional significance. New Phytologist, 164(2), 243–266.

Hawkins, H.-J., Cargill, R. I. M., Van Nuland, M. E., Hagen, S. C., Field, K. J., Sheldrake, M., Soudzilovskaia, N. A., & Kiers, E. T. (2023). Mycorrhizal mycelium as a global carbon pool. Current Biology, 33, R560–R573. 10.1016/j.cub.2023.02.027

Helfield, J. M., & Naiman, R. J. (2002). Salmon and alder as nitrogen sources to riparian forests in a boreal Alaskan watershed. Oecologia, 133, 573–582.

Hobbie, E. A., & Agerer, R. (2010). Nitrogen isotopes in ectomycorrhizal sporocarps correspond to belowground exploration types. Plant and Soil, 327, 71–83.

Hobbie, E. A., van Diepen, L. T. A., Lilleskov, E. A., Ouimette, A. P., Finzi, A. C., & Hofmockel, K. S. (2014). Fungal functioning in a pine forest: Evidence from a 15N-labeled global change experiment. New Phytologist, 201(4), 1431–1439.

Hobbie, E. A., Chen, J., & Hasselquist, N. J. (2019). Fertilization alters nitrogen isotopes and concentrations in ectomycorrhizal fungi and soil in pine forests. Fungal Ecology, 39, 267–275.

Hocking, M. D., & Reynolds, J. D. (2011). Impacts of salmon on riparian plant diversity. Science, 331(6024), 1609–1612.

Hocking, M. D., & Reynolds, J. D. (2012). Nitrogen uptake by plants subsidized by Pacific salmon carcasses: A hierarchical experiment. Canadian Journal of Forest Research, 42(5), 908–917.

Holtgrieve, G. W., Schindler, D. E., Gowell, C. P., Ruff, C. P., & Lisi, P. J. (2010). Stream geomorphology regulates the effects on periphyton of ecosystem engineering and nutrient enrichment by Pacific salmon. Freshwater Biology, 55, 2598–2611.

Hurteau, L. A., Mooers, A. Ø., Reynolds, J. D., & Hocking, M. D. (2016). Salmon nutrients are associated with the phylogenetic dispersion of riparian flowering-plant assemblages. Ecology, 97(2), 450–460.

Jach-Smith, L. C., & Jackson, R. D. (2020). Inorganic N addition replaces N supplied to switchgrass (*Panicum virgatum*) by arbuscular mycorrhizal fungi. Ecological Applications, 30(2), e02047.

Keski-Saari, S., & Julkunen-Tiitto, R. (2003). Resource allocation in different parts of juvenile mountain birch plants: Effect of nitrogen supply on seedling phenolics and growth. Physiologia Plantarum, 118(1), 114–126.

Kirchhoff, M. D. (2003). Effects of salmon-derived nitrogen on riparian forest growth and implications for stream productivity: Comment. Ecology, 84, 3396–3399.

Klock, A. M., Vogt, K. A., Vogt, D. J., Gordon, J. G., Scullion, J. J., Suntana, A. S., Mafune, K. K., Polyakov, A. Y., Gmur, S. J., & Gómez de la Rosa, C. (2022). See the forest not the trees! Ecosystem-based assessment of response, resilience, and scope for growth of global forests. Ecological Indicators, 140, 108973.

Kutorga, E., Irsenaite, R., Iznova, T., Kasparavicius, J., Markovskaja, S., & Motiejunaite, J. (2013). Species diversity and composition of fungal communities in a Scots pine forest affected by the great cormorant colony. Acta Mycologica, 48(2), 171–185.

Larocque, A. T. (2022). Fish, forests, fungi: Soils in the “salmon forests” of British Columbia (Doctoral dissertation, University of British Columbia).

Larocque, A. & Simard, S. W. (2023). Legacy of salmon-derived nutrients on riparian soil chemistry and soil fertility on the Central Coast of British Columbia, Canada. Frontiers in Forests and Global Change 6, 1010294.

Lilleskov, E. A., Hobbie, E. A., & Horton, T. R. (2011). Conservation of ectomycorrhizal fungi: Exploring the linkages between functional and taxonomic responses to anthropogenic N deposition. Fungal Ecology, 4(2), 174–183.

Lilleskov, E. A., Kuyper, T. W., Bidartondo, M. I., & Hobbie, E. A. (2019). Atmospheric nitrogen deposition impacts on the structure and function of forest mycorrhizal communities: A review. Environmental Pollution, 246, 148–162.

Lilleskov, E. A., Kuyper, T. W., Bidartondo, M. I. & Hobbie, E. A. (2024). Impacts of nitrogen manipulation on forest mycorrhizal communities in Atmospheric Nitrogen Manipulation to Global Forests. Academic Press, 2024, pp. 95–118.

Lodge, D. J., McDowell, W. H., & McSwiney, C. P. (1994). The importance of nutrient pulses in tropical forests. Trends in Ecology & Evolution, 9(10), 384–387.

Martin, M. (2011). Cutadapt removes adapter sequences from high-throughput sequencing reads. EMBnet.journal, 17(1), 10–12.

McMurdie, P. J., & Holmes, S. (2013). phyloseq: An R package for reproducible interactive analysis and graphics of microbiome census data. PLoS ONE, 8(4), e61217. 10.1371/journal.pone.0061217

Minakawa, N., Gara, R. I., & Honea, J. M. (2002). Increased individual growth rate and community biomass of stream insects associated with salmon carcasses. Journal of the North American Benthological Society, 21, 651–659.

Moeller, H. V., Peay, K. G., & Fukami, T. (2014). Ectomycorrhizal fungal traits reflect environmental conditions along a coastal California edaphic gradient. FEMS Microbiology Ecology, 87(3), 797–806.

Moeller, H. V., & Neubert, M. G. (2016). Multiple friends with benefits: An optimal mutualist management strategy? The American Naturalist, 187(1), E1–E12.

Nara, K. (2015). The role of ectomycorrhizal networks in seedling establishment and primary succession. In T. R. Horton (Ed.), Mycorrhizal networks (pp. 177–201). Springer.

Nguyen, N. H., Song, Z., Bates, S. T., Branco, S., Tedersoo, L., Menke, J., Schilling, J. S., & Kennedy, P. G. (2016). FUNGuild: An open annotation tool for parsing fungal community datasets by ecological guild. Fungal Ecology, 20, 241–248.

Nilsson, R. H., Larsson, K.-H., Taylor, A. F. S., Bengtsson-Palme, J., Jeppesen, T. S., Schigel, D., Kennedy, P., et al. (2019). The UNITE database for molecular identification of fungi: Handling dark taxa and parallel taxonomic classifications. Nucleic Acids Research, 47(D1), D259–D264.

Northrup, R. R., Yu, Z., Dahlgren, R. A., & Vogt, K. A. (1995). Polyphenol control of nitrogen release from pine litter. Nature, 377, 227–229.

Oksanen, J. (2016). Design decisions and implementation details in vegan (R package vignette, version 2–4).

Plassard, C., & Dell, B. (2010). Phosphorus nutrition of mycorrhizal trees. Tree Physiology, 30(9), 1129–1139.

Quinn, T. P., Helfield, J. M., Austin, C. S., Hovel, R. A., & Bunn, A. G. (2018). A multidecade experiment shows that fertilization by salmon carcasses enhanced tree growth in the riparian zone. Ecology, 99(11), 2433–2441.

Read, D. J., Leake, J. R., & Perez-Moreno, J. (2004). Mycorrhizal fungi as drivers of ecosystem processes in heathland and boreal forest biomes. Canadian Journal of Botany, 82(8), 1243–1263.

Rochet, J., Moreau, P.-A., Manzi, S., & Gardes, M. (2011). Comparative phylogenies and host specialization in the alder ectomycorrhizal fungi *Alnicola*, *Alpova*, and *Lactarius* (Basidiomycota) in Europe. BMC Evolutionary Biology, 11, 40.

Sagara, N. (1976). Presence of a buried mammalian carcass indicated by fungal fruiting bodies. Nature, 262(5571), 816.

Sagara, N., Yamanaka, T., & Tibbett, M. (2008). Soil fungi associated with graves and latrines: Toward a forensic mycology. In Soil analysis in forensic taphonomy (pp. 79–120). CRC Press.

Schindler, D. E., Armstrong, J. B., Bentley, K. T., Jankowski, K. J., Lisi, P. J., & Payne, L. X. (2013). Riding the crimson tide: Mobile terrestrial consumers track phenological variation in spawning of an anadromous fish. Biology Letters, 9(3), 20130048.

Siemens, L. D., Dennert, A. M., Obrist, D. S., & Reynolds, J. D. (2020). Spawning salmon density influences fruit production of salmonberry (*Rubus spectabilis*). Ecosphere, 11(11), e03282.

Treseder, K. K. (2008). Nitrogen additions and microbial biomass: A meta-analysis of ecosystem studies. Ecology Letters, 11(10), 1111–1120.

United States Department of Agriculture. (2022). Official soil series descriptions and series classifications. https://soilseries.sc.egov.usda.gov/OSD_Docs/A/ALEKNAGIK.html

Wagner, M. A., & Reynolds, J. D. (2019). Salmon increase forest bird abundance and diversity. PLoS ONE, 14(2), e0210031.

Walker, J. K. M., Cohen, H., Higgins, L. M., & Kennedy, P. G. (2014). Testing the link between community structure and function for ectomycorrhizal fungi involved in a global tripartite symbiosis. New Phytologist, 202(1), 287–296.

Wang, Q., Garrity, G. M., Tiedje, J. M., & Cole, J. R. (2007). Naive Bayesian classifier for rapid assignment of rRNA sequences into the new bacterial taxonomy. Applied and Environmental Microbiology, 73(16), 5261–5267.

Wang, J. R., Hawkins, C. D. B., & Letchford, T. (1998). Photosynthesis, water and nitrogen use efficiencies of four paper birch (*Betula papyrifera*) populations grown under different soil moisture and nutrient regimes. Forest Ecology and Management, 112(3), 233–244.

Weber, M. J., & Brown, M. L. (2013). Continuous, pulsed and disrupted nutrient subsidy effects on ecosystem productivity, stability, and energy flow. Ecosphere, 4(2), art27. 10.1890/ES12-00354.1

White, T. J., Bruns, T., Lee, S., & Taylor, J. W. (1990). Amplification and direct sequencing of fungal ribosomal RNA genes for phylogenetics. In M. A. Innis et al. (Eds.), PCR protocols: A guide to methods and applications (pp. 315–322). Academic Press.

Zanne, A. E., Abarenkov, K., Afkhami, M. E., Aguilar-Trigueros, C. A., Bates, S., Bhatnagar, J. M., Busby, P. E., et al. (2020). Fungal functional ecology: Bringing a trait-based approach to plant-associated fungi. Biological Reviews, 95(2), 409–433.

Zhao, Z.-B., He, J.-Z., Quan, Z., Wu, C.-F., Sheng, R., Zhang, L.-M., & Geisen, S. (2020). Fertilization changes soil microbiome functioning, especially phagotrophic protists. Soil Biology and Biochemistry, 148, 107863.

