## Supplementary Information for "Spatiotemporal differences in salmon nutrient inputs restructure functional and taxonomic fungal communities in riparian system"

**Supplementary Methods**

From 1997 to 2018, daily stream surveys were conducted during the entirety of the annual sockeye salmon run (approximately July 20 – August 20). During these surveys, all dead salmon in the creek and on the right stream bank up to 5 m were thrown onto the left stream bank. In addition, all salmon on the left bank were moved to a distance of about 3-6 m to avoid double-counting. As a result, almost all salmon carcasses at Hansen Creek were located roughly 3-6 m from the left bank of the stream, except for those relocated by wildlife and the salmon present following the end of stream surveys. This led to a decrease in salmon carcass density on the right side of the stream and an increase in salmon carcass density on the left side. Before the manipulation began in 1997, each bank of Hansen Creek received an average of 4,500 kg of salmon annually over an area of 1.2 ha (assuming an area 6 m wide by 2 km long), while after the manipulation began, the left bank received almost 10 times more salmon carcasses than the right bank (from the right bank and from the stream itself), averaging 13,400 kg of salmon while the right bank averaged 2,300 kg of salmon annually (due to some live salmon still present on the last survey date, which were assigned evenly between both banks; Quinn et al. 2018). Over the entire 20-year period, about 268,000 kg of salmon were moved to the left bank of Hansen Creek, equating to 8,000 kg of N and 1,400 kg of P, or 317 kg N/ha/year and 56 P/ha/year in a 6 m wide strip along the stream (Quinn et al. 2018, Feddern et al. 2019). After the cessation of the manipulations natural rates of salmon input were re-established.

(a)

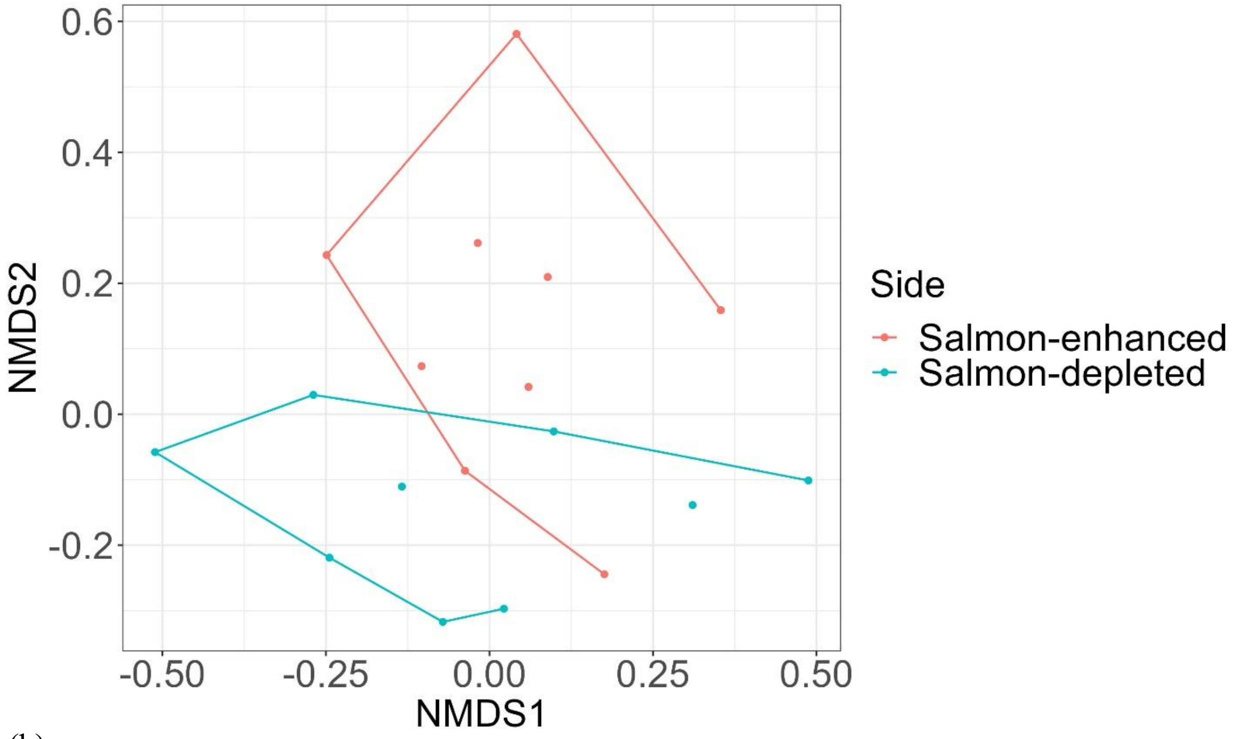


(b)

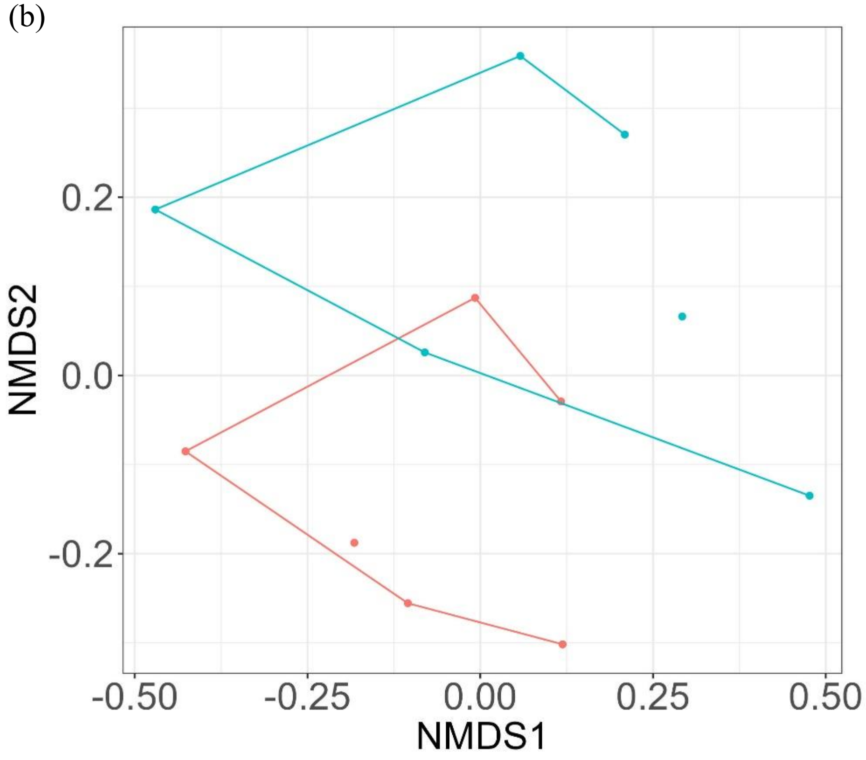


Figure S1. NMDS plot with lines connecting the outermost points for each group visualizing the effect of the salmon carcass deposition (on the salmon-enhanced bank) on fungal communities for (a) 1-6 m and (b) 3-6 m at Hansen Creek.

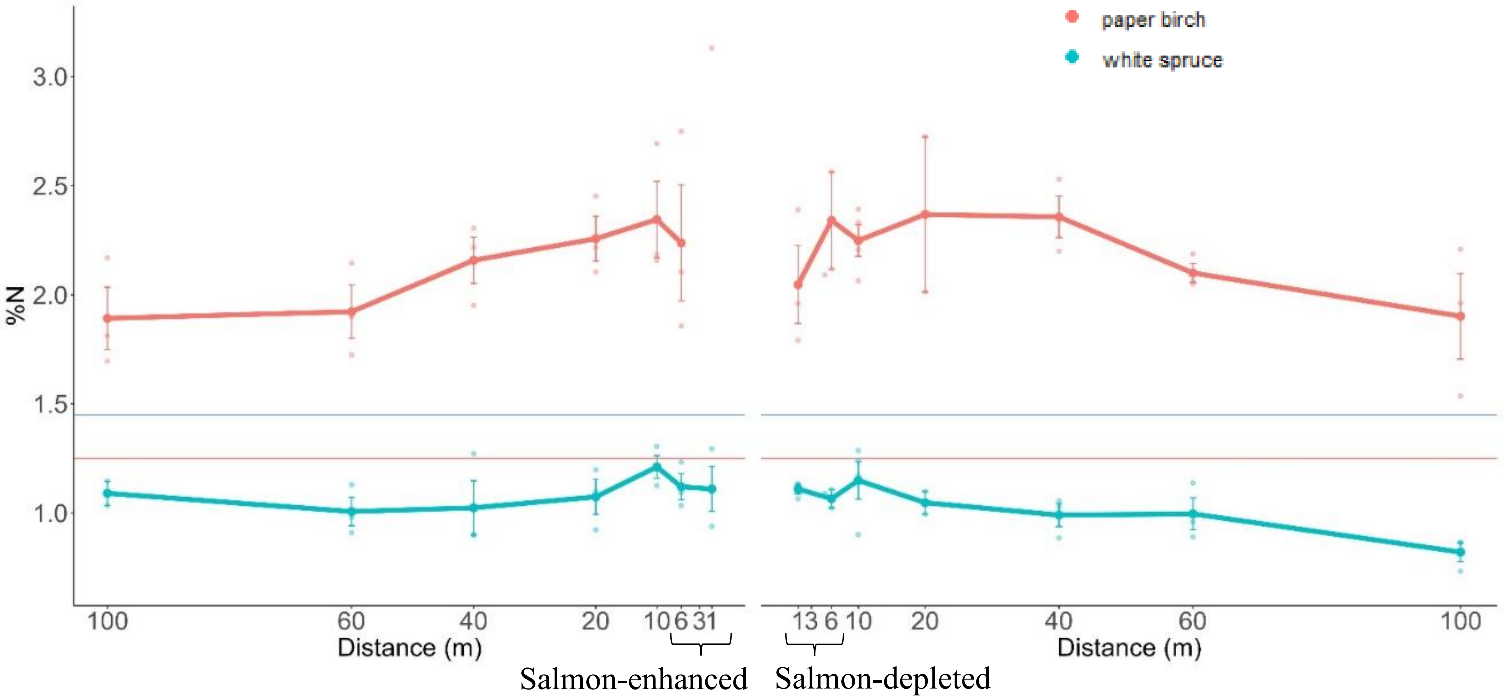


Figure S2. Mean values of foliar %N for white spruce and paper birch from 1-100 m away from the bank on both sides of Hansen Creek. Horizontal lines indicate values of N deficiency for white spruce (blue line) and paper birch (red line).

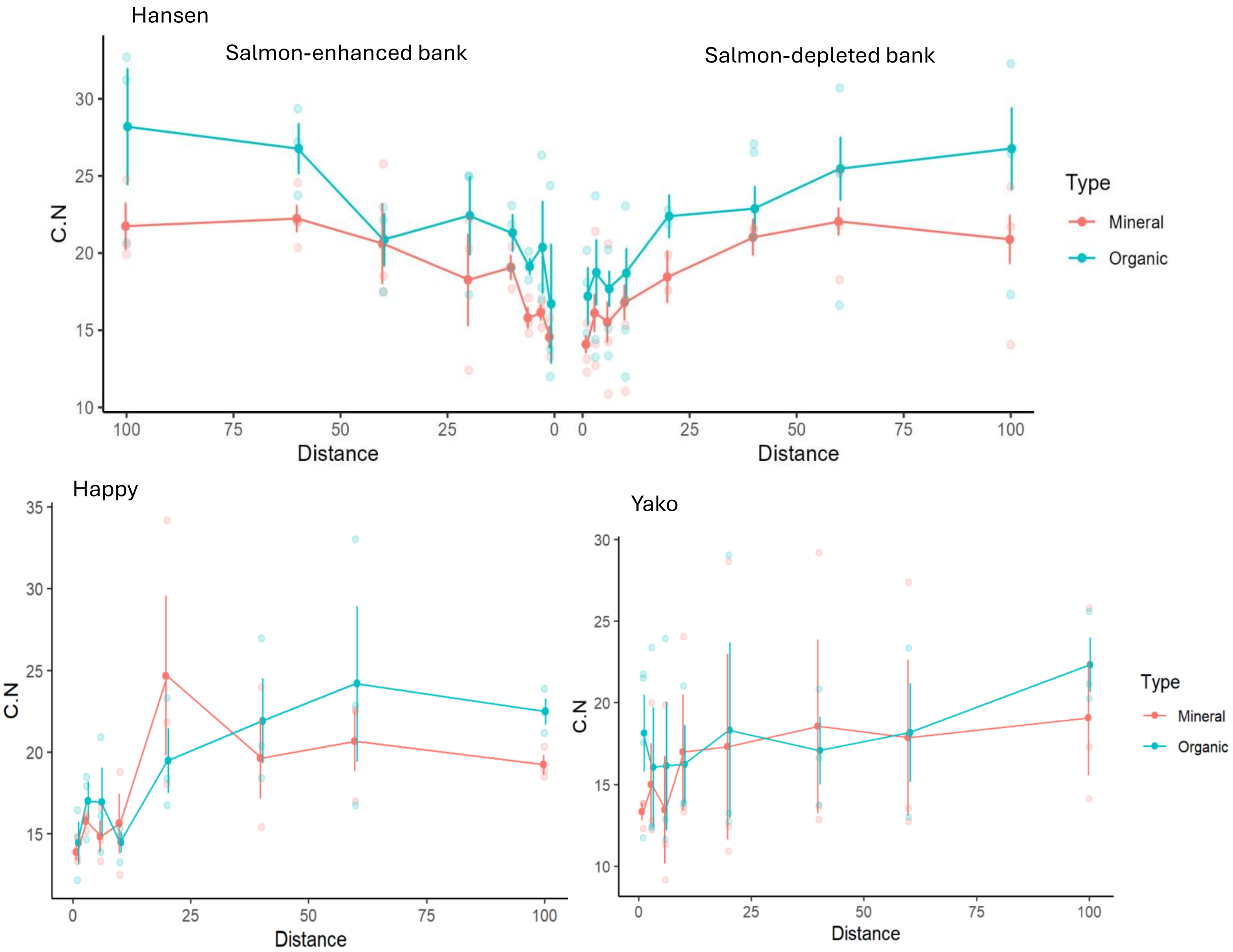


Figure S3. Organic and mineral soil C:N from 1-100 m away from both banks of Hansen Creek, and from 1-100 m away from the banks of Happy and Yako Creek.

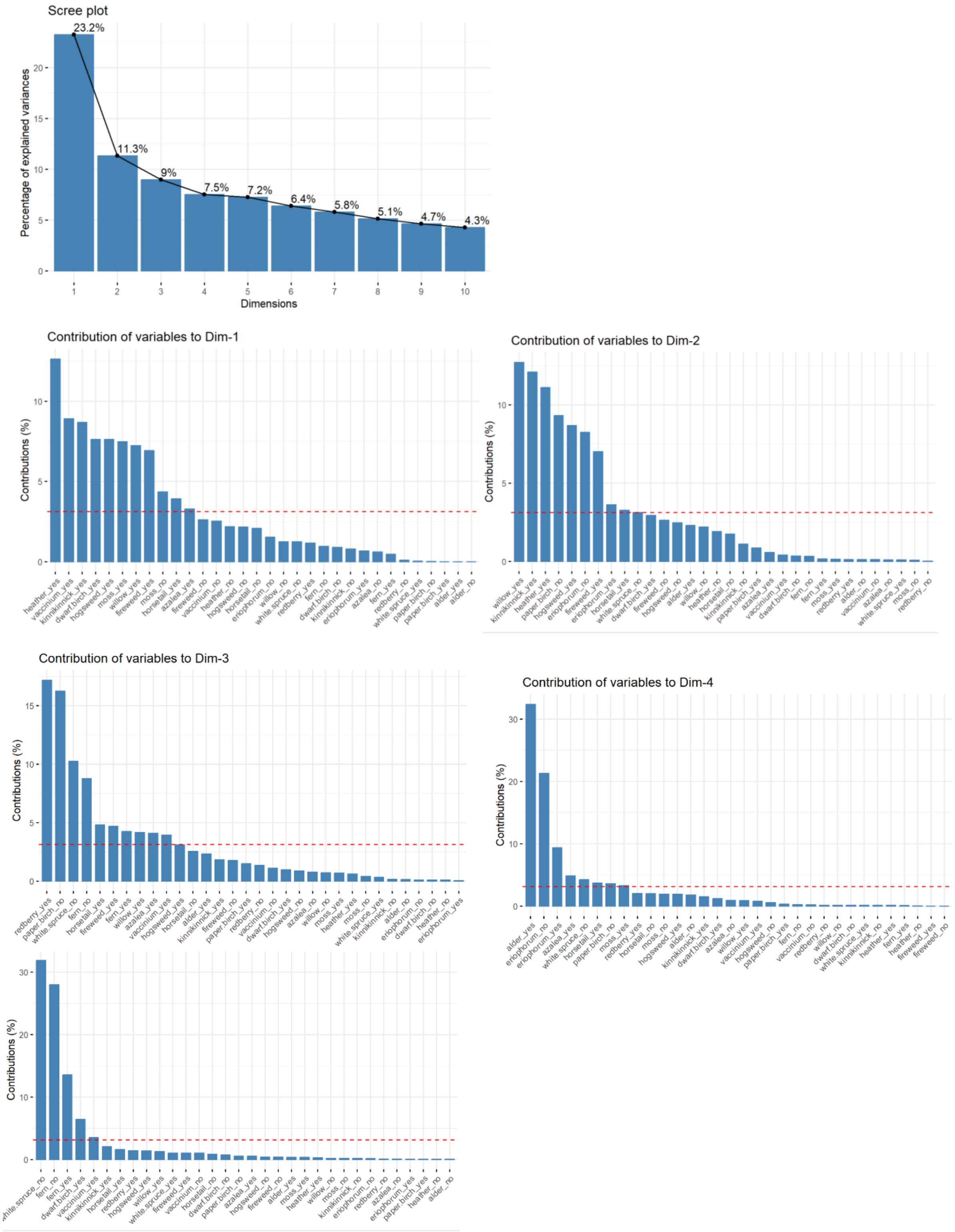


Figure S4. Plot of dimensions resulting from multiple correspondence analysis of plant community composition in a 5 $m^2$ radius surrounding soil sampling locations.

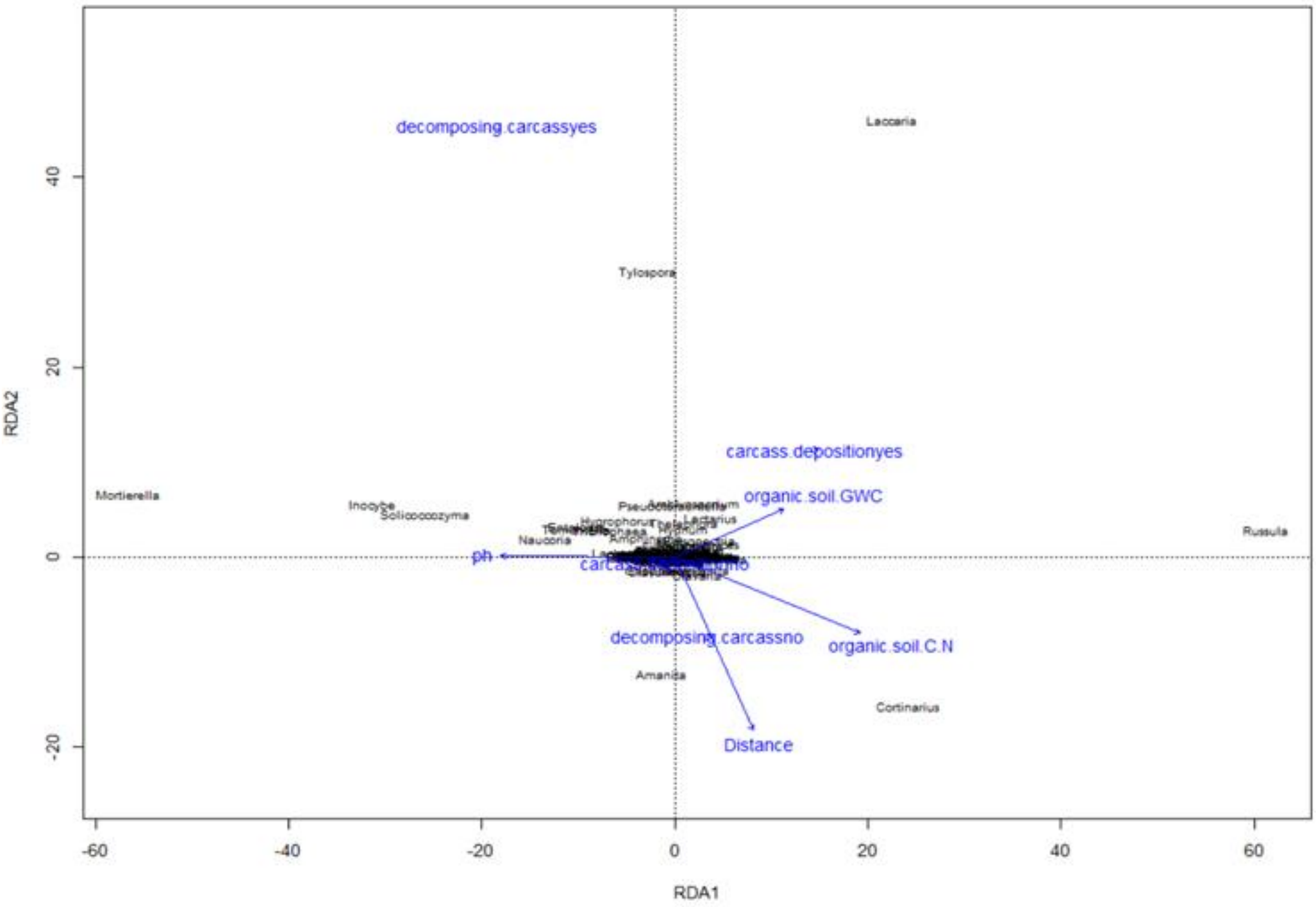


Figure S5. A correlation biplot representing redundancy analysis using 308 fungal genera and 6 environmental variables.

Table S1. Fungal species that were found only at sites near decomposing carcasses at three salmon streams in SW Alaska, ordered by trophic guild.

| Genus | Species | Trophic Guild | Type |
|---|---|---|---|
| *Metarhizium* | *carneum* | animal_parasite | none |
| *Beauveria* | *pseudobassiana* | animal_parasite | none |
| *Exophiala* | *bonariae* | animal_parasite | none |
| *Coprinus* | *phaeopunctatus* | dung_saprotroph | none |
| *Tomentella* | *viridula* | ectomycorrhizal | medium-distance_smooth |
| *Thelephora* | *palmata* | ectomycorrhizal | medium-distance_smooth |
| *Inocybe* | *mixtilis* | ectomycorrhizal | short-distance_delicate |
| *Cortinarius* | *annae-maritae* | ectomycorrhizal | medium-distance_fringe |
| *Hygrophorus* | *discoideus* | ectomycorrhizal | contact |
| *Inocybe* | *pseudodestricta* | ectomycorrhizal | short-distance_delicate |
| *Tomentella* | *clavigera* | ectomycorrhizal | medium-distance_smooth |
| *Otidea* | *nannfeldtii* | ectomycorrhizal | short-distance_coarse |
| *Cortinarius* | *glandicolor* | ectomycorrhizal | medium-distance_fringe |
| *Russula* | *helodes* | ectomycorrhizal | contact |
| *Cortinarius* | *pluvius* | ectomycorrhizal | medium-distance_fringe |
| *Cortinarius* | *disjungendulus* | ectomycorrhizal | medium-distance_fringe |
| *Alnicola* | *spectabilis* | ectomycorrhizal | short-distance_delicate |
| *Pseudotomentella* | *alnophila* | ectomycorrhizal | medium-distance_smooth |
| *Cortinarius* | *pluviorum* | ectomycorrhizal | medium-distance_fringe |
| *Cortinarius* | *pseudoturmalis* | ectomycorrhizal | medium-distance_fringe |
| *Cortinarius* | *acetosus* | ectomycorrhizal | medium-distance_fringe |
| *Cortinarius* | *illuminoides* | ectomycorrhizal | medium-distance_fringe |
| *Triposporium* | *cycadicola* | foliar_endophyte | none |
| *Monodictys* | *arctica* | lichen_parasite | none |
| *Psilocybe* | *medullosa* | litter_saprotroph | none |
| *Pyxidiophora* | *arvernensis* | litter_saprotroph | none |
| *Cystoderma* | *carpaticum* | litter_saprotroph | none |
| *Crocicreas* | *cyathoideum* | litter_saprotroph | none |
| *Deconica* | *merdaria* | litter_saprotroph | none |
| *Pluteus* | *leoninus* | litter_saprotroph | none |
| *Lepiota* | *magnispora* | litter_saprotroph | none |
| *Cystofilobasidium* | *capitatum* | litter_saprotroph | none |
| *Mycena* | *metata* | litter_saprotroph | none |
| *Neosetophoma* | *samararum* | litter_saprotroph | none |
| *Arrhenia* | *acerosa* | litter_saprotroph | none |
| *Deconica* | *bayliasiana* | litter_saprotroph | none |
| *Pluteus* | *eos* | litter_saprotroph | none |
| *Colacogloea* | *philyla* | mycoparasite | none |
| *Debaryomyces* | *hansenii* | nectar/tap_saprotroph | none |
| *Typhula* | *micans* | plant_pathogen | none |
| *Ilyonectria* | *mors-panacis* | plant_pathogen | none |

| | | | |
|---|---|---|---|
| *Polyscytalum* | *neofecundissimum* | plant_pathogen | none |
| *Microcyclospora* | *tardicrescens* | plant_pathogen | none |
| *Mortierella* | *sclerotiella* | soil_saprotroph | none |
| *Mortierella* | *rishikesha* | soil_saprotroph | none |
| *Phialocephala* | *virens* | soil_saprotroph | none |
| *Entoloma* | *nitens* | soil_saprotroph | none |
| *Coprinopsis* | *clastophylla* | soil_saprotroph | none |
| *Entoloma* | *calobrunneum* | soil_saprotroph | none |
| *Coprinopsis* | *romagnesiana* | soil_saprotroph | none |
| *Agaricus* | *macrocarpus* | soil_saprotroph | none |
| *Capronia* | *pulcherrima* | soil_saprotroph | none |
| *Leucosporidium* | *fellii* | soil_saprotroph | none |
| *Blastocladiella* | *britannica* | soil_saprotroph | none |
| *Entoloma* | *cuspidiferum* | soil_saprotroph | none |
| *Entoloma* | *rhombisporum* | soil_saprotroph | none |
| *Entoloma* | *vernum* | soil_saprotroph | none |
| *Solicoccozyma* | *terrea* | soil_saprotroph | none |
| *Entoloma* | *jubatum* | soil_saprotroph | none |
| *Entoloma* | *hirtipes* | soil_saprotroph | none |
| *Cuphophyllus* | *cinerellus* | soil_saprotroph | none |
| *Mrakia* | *niccombsii* | unspecified_saprotroph | none |
| *Claussenomyces* | *prasinulus* | wood_saprotroph | none |
| *Psathyrella* | *rostellata* | wood_saprotroph | none |
| *Lachnellula* | *calyciformis* | wood_saprotroph | none |
| *Coniophora* | *puteana* | wood_saprotroph | none |
| *Kuehneromyces* | *mutabilis* | wood_saprotroph | none |
| *Hyphodontiella* | *multiseptata* | wood_saprotroph | none |
| *Psathyrella* | *boreifasciculata* | wood_saprotroph | none |

Table S2. Predictors from the linear mixed effect models for relative abundance of each ectomycorrhizal fungal genus along three salmon streams in SW Alaska. Models include beta coefficient estimates ± SE, and P-values. Dim1 through Dim5 refer to the plant community metric vectors defined through multiple correspondence analysis (Figure S4). All models included a random effect of transect nested within stream. Organic soil GWC indicates organic soil gravimetric water content.

| **Long distance EMF** | | | | |
|---|---|---|---|---|
| **Genus** | **term** | **estimate** | **std.error** | **P value** |
| *Paxillus* | (Intercept) | -0.00343 | 0.002499 | 0.173201 |
| | Dim3 | -0.00308 | 0.000997 | 0.002671 |
| | pH | 0.001002 | 0.000592 | 0.093636 |
| | decomposing.carcassyes | 0.002241 | 0.000846 | 0.009512 |
| *Boletus* | (Intercept) | -5E-05 | 1.71E-05 | 0.004448 |
| | organic.soil.GWC | 6.29E-06 | 1.44E-06 | 3.49E-05 |
| | pH | 8.81E-06 | 3.9E-06 | 0.026156 |
| | carcass.depositionyes | 1.41E-05 | 7.29E-06 | 0.056532 |
| *Alpova* | (Intercept) | -0.01112 | 0.003356 | 0.001318 |
| | Dim3 | -0.0043 | 0.001363 | 0.002174 |
| | pH | 0.002901 | 0.000785 | 0.000375 |
| *Xerocomus* | (Intercept) | -0.00158 | 0.001031 | 0.12894 |
| | organic.soil.C.N | 0.000134 | 5.48E-05 | 0.01671 |
| | organic.soil.GWC | -0.00031 | 0.000171 | 0.070623 |
| | slopeyes | -0.00108 | 0.000697 | 0.125802 |
| | decomposing.carcassyes | 0.001514 | 0.000701 | 0.033365 |

| **Medium distance fringe EMF** | | | | |
|---|---|---|---|---|
| **Genus** | **term** | **estimate** | **std.error** | **p.value** |
| *Cortinarius* | (Intercept) | 0.00871 | 0.030503 | 0.775885 |
| | organic.soil.C.N | 0.006641 | 0.001807 | 0.000405 |
| | mineral.soil.C.N | -0.00545 | 0.002155 | 0.013213 |
| | slopeyes | 0.078337 | 0.020589 | 0.000258 |
| | carcass.depositionyes | -0.04496 | 0.022849 | 0.052161 |
| *Amphinema* | (Intercept) | 0.018615 | 0.005441 | 0.000939 |
| | mineral.soil.C.N | -0.00072 | 0.000291 | 0.015161 |
| | carcass.load | -4.7E-06 | 2.65E-06 | 0.081939 |
| | Dim3 | -0.0103 | 0.00454 | 0.025653 |
| | Dim5 | -0.00727 | 0.004977 | 0.147558 |
| *Piloderma* | (Intercept) | 0.001527 | 0.001645 | 0.355468 |
| | Dim3 | 0.00941 | 0.005147 | 0.070757 |
| | decomposing.carcassyes | 0.008582 | 0.004305 | 0.049191 |
| *Sistotrema* | (Intercept) | 5.97E-05 | 4.45E-05 | 0.183257 |
| | Dim1 | -0.00013 | 9.16E-05 | 0.155359 |
| | Dim2 | 0.00024 | 0.000129 | 0.065484 |

| | | | | |
|---|---|---|---|---|
| | Dim5 | -0.00023 | 0.000148 | 0.122327 |
| | decomposing.carcassyes | 0.000308 | 0.000131 | 0.02131 |
| *Tricholoma* | (Intercept) | 6.56E-05 | 3.26E-05 | 0.047048 |
| | Dim3 | 0.000369 | 0.00011 | 0.001142 |
| *Lyophyllum* | (Intercept) | 1.24E-05 | 7.64E-06 | 0.106739 |
| | Dim2 | 5.03E-05 | 2.29E-05 | 0.030608 |
| | Dim3 | -5.8E-05 | 2.58E-05 | 0.025886 |

| Medium distance smooth EMF | | | | |
|---|---|---|---|---|
| **Genus** | **term** | **estimate** | **std.error** | **p.value** |
| *Lactarius* | (Intercept) | 0.040211 | 0.008223 | 4.2E-06 |
| | organic.soil.GWC | -0.00611 | 0.003156 | 0.056082 |
| *Tomentella* | (Intercept) | 0.017052 | 0.004972 | 0.000907 |
| | Dim1 | -0.02979 | 0.009922 | 0.003443 |
| | carcass.depositionyes | 0.023465 | 0.016153 | 0.149725 |
| *Entoloma* | (Intercept) | 0.030067 | 0.008584 | 0.000722 |
| | ln(Distance) | -0.0057 | 0.00265 | 0.034128 |
| | Dim1 | -0.01947 | 0.007259 | 0.008717 |
| | Dim2 | 0.027476 | 0.009432 | 0.004526 |
| | decomposing.carcassyes | -0.01429 | 0.010297 | 0.168759 |
| | carcass.depositionyes | -0.01691 | 0.011439 | 0.142899 |
| *Pseudotomentella* | (Intercept) | 0.006971 | 0.002338 | 0.003665 |
| | Dim5 | -0.01546 | 0.008674 | 0.077937 |
| *Hydnum* | (Intercept) | -0.00712 | 0.010415 | 0.495881 |
| | organic.soil.C.N | 0.000829 | 0.000529 | 0.120352 |
| | ln(Distance) | -0.00355 | 0.001976 | 0.075737 |
| | Dim3 | 0.01413 | 0.008428 | 0.097097 |
| | decomposing.carcassyes | 0.016035 | 0.007478 | 0.034712 |
| *Thelephora* | (Intercept) | 0.005351 | 0.003068 | 0.084373 |
| | carcass.depositionyes | 0.019619 | 0.009966 | 0.051986 |
| *Tomentellopsis* | (Intercept) | -0.00055 | 0.000223 | 0.01512 |
| | organic.soil.C.N | -1.7E-05 | 1.1E-05 | 0.130362 |
| | mineral.soil.C.N | 5.51E-05 | 1.24E-05 | 2.37E-05 |
| | slopeyes | -0.0002 | 0.000116 | 0.082141 |
| | Dim1 | -0.00023 | 0.000113 | 0.043887 |

| Short distance EMF | | | | |
|---|---|---|---|---|
| **Genus** | **term** | **estimate** | **std.error** | **p.value** |
| *Russula* | (Intercept) | 0.413231 | 0.194111 | 0.035969 |
| | Dim1 | 0.092489 | 0.048111 | 0.057681 |
| | Dim3 | 0.137644 | 0.067414 | 0.044069 |
| | pH | -0.064 | 0.045548 | 0.163362 |
| *Naucoria* | (Intercept) | -0.17325 | 0.045458 | 0.00025 |

| | | | | |
|---|---|---|---|---|
| | organic.soil.GWC | 0.00806 | 0.003601 | 0.027593 |
| | pH | 0.040182 | 0.010365 | 0.000198 |
| *Inocybe* | (Intercept) | 0.081947 | 0.013727 | 4.41E-08 |
| | Dim5 | 0.082717 | 0.048493 | 0.091428 |
| | carcass.depositionyes | -0.07819 | 0.044634 | 0.083155 |
| *Clavulina* | (Intercept) | -0.01442 | 0.007338 | 0.052553 |
| | organic.soil.GWC | 0.001016 | 0.000616 | 0.102483 |
| | carcass.load | 4.06E-06 | 1.72E-06 | 0.020674 |
| | Dim3 | 0.004581 | 0.002944 | 0.123211 |
| | pH | 0.003353 | 0.001681 | 0.049086 |
| | carcass.depositionyes | -0.00631 | 0.003273 | 0.056921 |
| *Laccaria* | (Intercept) | 0.397047 | 0.112501 | 0.000653 |
| | ln(Distance) | -0.0268 | 0.008384 | 0.00191 |
| | pH | -0.06008 | 0.024131 | 0.014586 |
| *Tylospora* | (Intercept) | 0.179398 | 0.066568 | 0.008432 |
| | organic.soil.C.N | -0.01089 | 0.003219 | 0.001068 |
| | mineral.soil.C.N | 0.005056 | 0.003406 | 0.141236 |
| | carcass.load | 3.52E-05 | 2.14E-05 | 0.104168 |
| | Dim1 | 0.063893 | 0.032256 | 0.050735 |
| | Dim3 | 0.061346 | 0.036802 | 0.09909 |
| | carcass.depositionyes | 0.090083 | 0.037586 | 0.01866 |
| *Hebeloma* | (Intercept) | -0.00502 | 0.002056 | 0.016688 |
| | organic.soil.GWC | 0.000333 | 0.000153 | 0.031878 |
| | Dim1 | -0.00114 | 0.000561 | 0.045865 |
| | Dim2 | 0.001716 | 0.000722 | 0.019555 |
| | pH | 0.00118 | 0.000485 | 0.017008 |
| | decomposing.carcassyes | -0.00134 | 0.000682 | 0.052746 |
| *Meliniomyces* | (Intercept) | -0.00435 | 0.002168 | 0.04781 |
| | organic.soil.C.N | 0.000522 | 0.000122 | 4.8E-05 |
| | ln(Distance) | -0.00091 | 0.000461 | 0.051291 |
| | carcass.depositionyes | 0.004533 | 0.00204 | 0.028749 |
| *Tuber* | (Intercept) | -0.01021 | 0.002913 | 0.000719 |
| | mineral.soil.C.N | 0.000162 | 6.18E-05 | 0.010137 |
| | organic.soil.GWC | -0.00033 | 0.000149 | 0.029527 |
| | ln(Distance) | -0.00062 | 0.000181 | 0.000955 |
| | Dim3 | -0.00122 | 0.000775 | 0.118992 |
| | Dim5 | 0.004606 | 0.000826 | 2.64E-07 |
| | pH | 0.002438 | 0.000523 | 1.1E-05 |
| *Leotia* | (Intercept) | 0.006415 | 0.003228 | 0.049905 |
| | mineral.soil.C.N | -0.00013 | 6.73E-05 | 0.060028 |
| | ln(Distance) | 0.000463 | 0.000189 | 0.016053 |
| | Dim4 | 0.001537 | 0.000903 | 0.092286 |
| | pH | -0.00116 | 0.000579 | 0.047838 |
| *Sebacina* | (Intercept) | -0.00659 | 0.002976 | 0.029201 |

| | | | | |
|---|---|---|---|---|
| | organic.soil.C.N | 8.7E-05 | 5.41E-05 | 0.111365 |
| | organic.soil.GWC | -0.00027 | 0.000149 | 0.072397 |
| | Dim5 | -0.00116 | 0.00077 | 0.135333 |
| | pH | 0.001387 | 0.000526 | 0.00984 |
| *Alnicola* | (Intercept) | -0.00198 | 0.001062 | 0.066006 |
| | mineral.soil.C.N | 3.34E-05 | 2.11E-05 | 0.11591 |
| | organic.soil.GWC | -0.00016 | 5.87E-05 | 0.009548 |
| | carcass.load | 2.97E-07 | 1.57E-07 | 0.062577 |
| | Dim2 | -0.00039 | 0.000246 | 0.112132 |
| | pH | 0.000402 | 0.000188 | 0.034994 |
| | carcass.depositionyes | 0.000853 | 0.000298 | 0.005262 |
| *Hymenogaster* | (Intercept) | 0.001379 | 0.000879 | 0.120139 |
| | organic.soil.C.N | -8.2E-05 | 4.52E-05 | 0.073243 |
| | organic.soil.GWC | 0.000248 | 0.00016 | 0.125241 |
| *Pulvinula* | (Intercept) | -0.00076 | 0.000291 | 0.010704 |
| | Dim3 | 0.000255 | 0.000118 | 0.033319 |
| | Dim5 | -0.00023 | 0.000127 | 0.068934 |
| | pH | 0.000187 | 6.81E-05 | 0.007201 |
| *Endogone* | (Intercept) | -0.00076 | 0.000291 | 0.010704 |
| | Dim3 | 0.000255 | 0.000118 | 0.033319 |
| | Dim5 | -0.00023 | 0.000127 | 0.068934 |
| | pH | 0.000187 | 6.81E-05 | 0.007201 |
| *Trichophaea* | (Intercept) | 0.05578 | 0.010149 | 3.58E-07 |
| | organic.soil.C.N | -0.00099 | 0.000584 | 0.092916 |
| | mineral.soil.C.N | -0.0013 | 0.000662 | 0.053078 |
| | decomposing.carcassyes | -0.01369 | 0.006567 | 0.039895 |
| | carcass.depositionyes | -0.01144 | 0.007475 | 0.129512 |
| *Wilcoxina* | (Intercept) | 0.005395 | 0.003897 | 0.169613 |
| | organic.soil.C.N | -0.0006 | 0.000196 | 0.002783 |
| | mineral.soil.C.N | 0.000448 | 0.00021 | 0.035055 |
| | Dim1 | 0.004581 | 0.001995 | 0.023929 |
| *Cenococcum* | (Intercept) | -0.00711 | 0.003632 | 0.05329 |
| | organic.soil.C.N | 0.000412 | 0.000183 | 0.027282 |
| | slopeyes | 0.003975 | 0.002078 | 0.058915 |
| | Dim1 | -0.00572 | 0.002064 | 0.006786 |
| | Dim4 | -0.0071 | 0.002519 | 0.005936 |
| *Genabea* | (Intercept) | -0.00223 | 0.000854 | 0.010704 |
| | Dim3 | 0.000749 | 0.000347 | 0.033319 |
| | Dim5 | -0.00069 | 0.000373 | 0.068934 |
| | pH | 0.00055 | 0.0002 | 0.007201 |
| *Otidea* | (Intercept) | 5.47E-06 | 1.58E-05 | 0.730311 |
| | Dim2 | 9.86E-05 | 4.61E-05 | 0.035099 |
| | decomposing.carcassyes | 7.02E-05 | 4.34E-05 | 0.108929 |
| *Geopora* | (Intercept) | 1.32E-08 | 1.07E-06 | 0.990155 |

| | | | | |
|---|---|---|---|---|
| | Dim5 | -5.5E-06 | 3.65E-06 | 0.134006 |
| | decomposing.carcassyes | 6.95E-06 | 2.78E-06 | 0.014083 |
| | (Intercept) | 0.003625 | 0.010886 | 0.73988 |
| *Amanita* | organic.soil.GWC | -0.00622 | 0.003032 | 0.043071 |
| | slopeyes | -0.02022 | 0.012682 | 0.114307 |
| | ln(Distance) | 0.008358 | 0.003271 | 0.012286 |
| | Dim2 | -0.02425 | 0.013585 | 0.077588 |
| *Hygrophorus* | (Intercept) | 0.050716 | 0.011501 | 2.83E-05 |
| | organic.soil.C.N | -0.0015 | 0.00058 | 0.011345 |
| | mineral.soil.C.N | -0.00089 | 0.000618 | 0.155464 |
| | Dim1 | 0.014642 | 0.005887 | 0.014693 |
| *Helvellosebacina* | (Intercept) | 8.58E-08 | 6.93E-06 | 0.990155 |
| | Dim5 | -3.6E-05 | 2.37E-05 | 0.134006 |
| | decomposing.carcassyes | 4.52E-05 | 1.81E-05 | 0.014083 |

Table S3. The %P, %N, and N:P values for paper birch, and white spruce samples at Hansen Creek at 1, 6, and 100 m on both the salmon-enhanced (SL) and salmon-depleted (SR) banks, in addition to four ectomycorrhizal fungal sporocarps and three organic soil samples.

| Stream | Transect | Side | Distance | Type | % P | % N | N:P |
|---|---|---|---|---|---|---|---|
| Hansen | S6 | SL | 6 | Lactarius tabidus | 0.38 | 5.02 | 13.13 |
| Hansen | S6 | SR | 1 | Lactarius tabidus | 0.53 | 4.09 | 7.74 |
| Hansen | S4 | SL | 4 | Paxillus involutus | 0.79 | 4.23 | 5.34 |
| Hansen | S1 | SR | 10 | Paxillus involutus | 0.69 | 4.80 | 6.91 |
| Hansen | S4 | SL | 1 | organic soil | 0.14 | 0.71 | 5.13 |
| Hansen | S4 | SL | 100 | organic soil | 0.08 | 1.30 | 16.16 |
| Hansen | S4 | SR | 1 | organic soil | 0.12 | 2.54 | 22.00 |
| Hansen | S4 | SL | 6 | paper birch | 0.17 | 2.11 | 12.69 |
| Hansen | S4 | SL | 100 | paper birch | 0.14 | 1.81 | 13.07 |
| Hansen | S4 | SR | 1 | paper birch | 0.22 | 2.39 | 10.78 |
| Hansen | S1 | SR | 1 | paper birch | 0.24 | 1.79 | 7.36 |
| Hansen | S6 | SR | 5 | paper birch | 0.22 | 2.09 | 9.35 |
| Hansen | S6 | SL | 6 | paper birch | 0.18 | 1.86 | 10.43 |
| Hansen | S1 | SL | 6 | paper birch | 0.42 | 2.75 | 6.49 |
| Hansen | S4 | SR | 6 | paper birch | 0.23 | 2.12 | 9.21 |
| Hansen | S4 | SR | 100 | paper birch | 0.32 | 2.21 | 6.94 |
| Hansen | S1 | SR | 6 | paper birch | 0.28 | 2.56 | 9.13 |
| Hansen | S6 | SR | 1 | paper birch | 0.15 | 1.96 | 13.04 |
| Hansen | S1 | SL | 1 | paper birch | 0.34 | 3.13 | 9.13 |
| Hansen | S4 | SL | 1 | white spruce | 0.14 | 1.29 | 9.10 |
| Hansen | S4 | SL | 100 | white spruce | 0.13 | 1.03 | 7.78 |
| Hansen | S4 | SR | 1 | white spruce | 0.13 | 1.13 | 8.65 |
| Hansen | S4 | SL | 6 | white spruce | 0.15 | 1.03 | 7.10 |
| Hansen | S1 | SL | 1 | white spruce | 0.15 | 0.94 | 6.42 |
| Hansen | S4 | SR | 6 | white spruce | 0.22 | 1.11 | 5.06 |
| Hansen | S4 | SR | 100 | white spruce | 0.15 | 0.87 | 5.58 |
| Hansen | S1 | SL | 6 | white spruce | 0.14 | 1.09 | 7.93 |
| Hansen | S6 | SR | 5 | white spruce | 0.15 | 1.09 | 7.34 |
| Hansen | S1 | SR | 1 | white spruce | 0.15 | 1.07 | 7.13 |
| Hansen | S1 | SR | 6 | white spruce | 0.13 | 1.02 | 7.98 |
| Hansen | S6 | SL | 6 | white spruce | 0.16 | 1.23 | 7.76 |
| Hansen | S6 | SL | 1 | white spruce | 0.16 | 1.10 | 6.78 |
| Hansen | S6 | SR | 1 | white spruce | 0.16 | 1.14 | 7.22 |

Table S4. List of all plant species observed within a 5 $m^2$ radius of soil sampling locations along transects perpendicular to three salmon streams in SW Alaska.

| Latin name | Common name |
|---|---|
| *Alnus crispa* | Green alder |
| *Arctostaphylos uva-ursi* | Kinnikinnick<br>Bearberry |
| *Betula nana* | Dwarf birch |
| *Betula papyrifera* | Paper birch |
| *Chamaenerion angustifolium* | Fireweed |
| *Equisetum arvense* | Horsetail |
| *Eriophorum* spp. | Cotton grass |
| *Harrimanella stelleriana* | Alaskan moss heather |
| Heracleum mantegazzianum | Hogsweed |
| *Picea glauca* | White spruce |
| *Pteridium aquilinum* | Bracken fern |
| *Salix alaxensis* | Felt leaf willow |
| *Sphagnum* spp. | Mosses |
| *Vaccinium ovalifolium* | Alaskan blueberry |
| *Vaccinium vitis-idaea* | Lowbush mountain lingonberry |

Table S5. Indicator species determined from species indicator analysis and primary lifestyles significantly associated with (a) the presence of nearby decomposing carcasses across three salmon streams in SW Alaska, and (b) the salmon-enhanced bank where salmon carcasses were deposited or the salmon-depleted bank where salmon carcasses were removed from 1-6 m at Hansen Creek during a 21-year salmon carcass manipulation experiment at Hansen Creek in SW Alaska.

(a)

| **Species** | **Lifestyle** |
|---|---|
| *Lophium arboricola* | litter saprotroph |
| *Trichocladium opacum* | unspecified saprotroph |
| *Russula consobrina* | ectomycorrhizal, short-distance type |
| *Hypholoma frowardii* | wood saprotroph |
| *Mortierella globulifera* | soil saprotroph |
| *Acrodontium hydnicola* | plant pathogen and litter saprotroph |
| *Hygrophorus olivaceoalbus* | ectomycorrhizal, short-distance type |
| *Lactarius rufus* | ectomycorrhizal, medium-distance type |
| *Mortierella sclerotiella* | soil saprotroph |
| *Cortinarius casimiri* | ectomycorrhizal, medium-distance fringe |

(b)

| **Genus** | **Lifestyle** | **Side** |
|---|---|---|
| *Ophiocordycipitacea inflatum* | Animal parasite/decomposer | Salmon-enhanced |
| *Oidiodendron trunca* | Soil saprotroph/root endophyte | Salmon-enhanced |
| *Amblyosporium botryt* | Mycoparasite/fungal decomposer | Salmon-enhanced |
| *Tomentella sublilacina* | ectomycorrhizal, medium smooth type | Salmon-enhanced |
| *Inocybe napipes* | ectomycorrhizal, short-distance type | Salmon-depleted |
| *Mortierella cystojenkinii* | soil saprotroph | Salmon-depleted |
| *Lactarius vietus* | ectomycorrhizal, medium-smooth type | Salmon-depleted |

Table S6. All ectomycorrhizal fungal species found along three salmon streams in SW Alaska, categorized by exploration types based on genus, where sp. refers to the lack of species-level ID for some samples.

| **Long distance species** |
|---|
| *Paxillus involutus* |
| *Alpova corsicus* |
| *Alpova sp.* |
| *Xerocomus sp.* |
| *Boletus edulis* |

| **Medium distance fringe species** |
|---|
| *Amphinema byssoides* |
| *Amphinema diadema* |
| *Amphinema sp.* |
| *Cortinarius acetosus* |
| *Cortinarius acutovelatus* |
| *Cortinarius acutus* |
| *Cortinarius adalbertii* |
| *Cortinarius alboamarescens* |
| *Cortinarius alnetorum* |
| *Cortinarius alpinus* |
| *Cortinarius annae-maritae* |
| *Cortinarius anomalus* |
| *Cortinarius armillatus* |
| *Cortinarius aurantiobasis* |
| *Cortinarius badiolatus* |
| *Cortinarius barlowensis* |
| *Cortinarius caninus* |
| *Cortinarius caperatus* |
| *Cortinarius casimiri* |
| *Cortinarius comptulus* |
| *Cortinarius delibutus* |
| *Cortinarius disjungendulus* |
| *Cortinarius duristipes* |
| *Cortinarius evernius* |
| *Cortinarius fennoscandicus* |
| *Cortinarius ferruginosus* |
| *Cortinarius fulvescens* |
| *Cortinarius fuscescens* |
| *Cortinarius glandicolor* |

| |
|---|
| *Cortinarius illibatus* |
| *Cortinarius illuminoides* |
| *Cortinarius ionophyllus* |
| *Cortinarius ionosmus* |
| *Cortinarius jonimitchelliae* |
| *Cortinarius leiocastaneus* |
| *Cortinarius lucorum* |
| *Cortinarius luteo-ornatus* |
| *Cortinarius malicorius* |
| *Cortinarius millaresensis* |
| *Cortinarius multiformis* |
| *Cortinarius sp.* |
| *Cortinarius obtusus* |
| *Cortinarius ochrophyllus* |
| *Cortinarius panellus* |
| *Cortinarius pluviorum* |
| *Cortinarius pluvius* |
| *Cortinarius pseudoturmalis* |
| *Cortinarius rufolatus* |
| *Cortinarius scaurus* |
| *Cortinarius septentrionalis* |
| *Cortinarius spilomeus* |
| *Cortinarius subpaleaceus* |
| *Cortinarius transatlanticus* |
| *Cortinarius valgus* |
| *Cortinarius var._notandus* |
| *Cortinarius vibratilis* |
| *Cortinarius violaceus* |
| *Cortinarius xanthocephalus* |
| *Lyophyllum sp.* |
| *Piloderma bicolor* |
| *Piloderma lanatum* |
| *Piloderma sphaerosporum* |
| *Sistotrema luteoviride* |
| *Sistotrema muscicola* |
| *Tricholoma fulvum* |
| *Tricholoma sp.* |
| *Tricholoma saponaceum* |
| *Tricholoma stiparophyllum* |

| **Medium distance smooth species** |
| --- |
| *Entoloma borgenii* |
| *Entoloma paludicola* |
| *Entoloma politum* |
| *Entoloma rhodopolium* |
| *Entoloma rubrobasis* |
| *Entoloma sericatum* |
| *Entoloma serpens* |
| *Hydnum sp.* |
| *Lactarius atroviridis* |
| *Lactarius flexuosus* |
| *Lactarius glyciosmus* |
| *Lactarius hengduanensis* |
| *Lactarius lapponicus* |
| *Lactarius sp.* |
| *Lactarius necator* |
| *Lactarius rufus* |
| *Lactarius tabidus* |
| *Lactarius tesquorum* |
| *Lactarius trivialis* |
| *Lactarius vietus* |
| *Pseudotomentella alnophila* |
| *Pseudotomentella mucidula* |
| *Pseudotomentella sp.* |
| *Pseudotomentella pluriloba* |
| *Pseudotomentella umbrina* |
| *Tomentella badia* |
| *Tomentella bryophila* |
| *Tomentella clavigera* |
| *Tomentella coerulea* |
| *Tomentella ellisii* |
| *Tomentella lapida* |
| *Tomentella lateritia* |
| *Tomentella sp.* |
| *Tomentella stuposa* |
| *Tomentella subclavigera* |
| *Tomentella sublilacina* |
| *Tomentella viridula* |
| *Tomentellopsis sp.* |
| *Tomentellopsis submollis* |

| **Short distance species** |
|---|
| *Alnicola cholea* |
| *Alnicola sp.* |
| *Alnicola spectabilis* |
| *Amanita constricta* |
| *Amanita magnivolvata* |
| *Amanita sp.* |
| *Ambispora sp.* |
| *Aspicilia sp.* |
| *Cadophora orchidicola* |
| *Cenococcum geophilum* |
| *Cenococcum sp.* |
| *Clavulina coralloides* |
| *Clavulina sp.* |
| *Clavulina ornatipes* |
| *Cryptodiscus sp.* |
| *Genabea sp.* |
| *Geopora sp.* |
| *Gymnopilus penetrans* |
| *Hebeloma aurantioumbrinum* |
| *Hebeloma cavipes* |
| *Hebeloma hetieri* |
| *Hebeloma hiemale* |
| *Hebeloma mesophaeum* |
| *Hebeloma sp.* |
| *Helvellosebacina sp.* |
| *Hygrophorus discoideus* |
| *Hygrophorus monticola* |
| *Hygrophorus olivaceoalbus* |
| *Hygrophorus pustulatus* |
| *Hymenogaster sp.* |
| *Hymenogaster rubyensis* |
| *Inocybe acuta* |
| *Inocybe ambigua* |
| *Inocybe assimilata* |
| *Inocybe borealis* |
| *Inocybe catalaunica* |
| *Inocybe flavella* |
| *Inocybe geophylla* |
| *Inocybe grammata* |
| *Inocybe impexa* |

| |
|---|
| *Inocybe lacera* |
| *Inocybe lanuginosa* |
| *Inocybe lapponica* |
| *Inocybe mixtilis* |
| *Inocybe sp.* |
| *Inocybe napipes* |
| *Inocybe nitidiuscula* |
| *Inocybe nothomixtilis* |
| *Inocybe pallidicremea* |
| *Inocybe posterula* |
| *Inocybe pseudodestricta* |
| *Inocybe rufoalba* |
| *Inocybe soluta* |
| *Inocybe stellatospora* |
| *Inocybe substellata* |
| *Inocybe teraturgus* |
| *Inocybe tetragonospora* |
| *Inocybe whitei* |
| *Laccaria laccata* |
| *Laccaria sp.* |
| *Laccaria pseudomontana* |
| *Naucoria bohemica* |
| *Naucoria sp.* |
| *Oidiodendron maius* |
| *Otidea nannfeldtii* |
| *Pulvinula sp.* |
| *Rhodoscypha sp.* |
| *Russula adusta* |
| *Russula amoenoides* |
| *Russula cessans* |
| *Russula citrinochlora* |
| *Russula claroflava* |
| *Russula consobrina* |
| *Russula cupreola* |
| *Russula decolorans* |
| *Russula emetica* |
| *Russula favrei* |
| *Russula griseascens* |
| *Russula helodes* |
| *Russula intermedia* |
| *Russula murrillii* |

| |
|---|
| *Russula sp.* |
| *Russula nuoljae* |
| *Russula paludosa* |
| *Russula sapinea* |
| *Russula suecica* |
| *Russula versicolor* |
| *Russula vinososordida* |
| *Russula violaceoincarnata* |
| *Russula viscida* |
| *Sebacina sp.* |
| *Tomentellopsis sp.* |
| *Tomentellopsis submollis* |
| *Trichophaea sp.* |
| *Tuber sp.* |
| *Tuber wenchuanense* |
| *Tylospora asterophora* |
| *Tylospora fibrillosa* |
| *Wilcoxina mikolae* |
